# GWAS of 19,629 individuals identifies novel genetic variants for regional brain volumes and refines their genetic co-architecture with cognitive and mental health traits

**DOI:** 10.1101/586339

**Authors:** Bingxin Zhao, Tianyou Luo, Tengfei Li, Yun Li, Jingwen Zhang, Yue Shan, Xifeng Wang, Liuqing Yang, Fan Zhou, Ziliang Zhu, Hongtu Zhu

**Affiliations:** Departments of Biostatistics, University of North Carolina at Chapel Hill, Chapel Hill, NC, USA, 27599; Departments of Radiology, University of North Carolina at Chapel Hill, Chapel Hill, NC, USA, 27599; Biomedical Research Imaging Center, School of Medicine, The University of North Carolina at Chapel Hill, Chapel Hill, NC, USA, 27599; Departments of Computer Science, University of North Carolina at Chapel Hill, Chapel Hill, NC, USA, 27599; Departments of Genetics, University of North Carolina at Chapel Hill, Chapel Hill, NC, USA, 27599; Department of Biostatistics, T.H. Chan School of Public Health, Harvard University, Boston, MA, USA, 02115; Departments of Statistics and Operations Research, University of North Carolina at Chapel Hill, Chapel Hill, NC, USA, 27599

**Keywords:** Genetic co-architecture, Genetic correlation, Pleiotropy, UK Biobank, Brain structure, Regional brain volumes.

## Abstract

Volumetric variations of human brain are heritable and are associated with many brain-related complex traits. Here we performed genome-wide association studies (GWAS) and post-GWAS analyses of 101 brain volumetric phenotypes using the UK Biobank (UKB) sample including 19,629 participants. GWAS identified 287 independent SNPs exceeding genome-wide significance threshold of 4.9*10^−10^, adjusted for testing multiple phenotypes. Gene-based association study found 142 associated genes (113 new) and functional gene mapping analysis linked 122 more genes. Many of the discovered genetic variants have previously been implicated with cognitive and mental health traits (such as cognitive performance, education, mental disease/disorders), and significant genetic correlations were detected for 29 pairs of traits. The significant SNPs discovered in the UKB sample were supported by a joint analysis with other four independent studies (total sample size 2,192), and we performed a meta-analysis of five samples to provide GWAS summary statistics with sample size larger than 20,000. Using genome-wide polygenic risk scores prediction, up to 4.36% of phenotypic variance (p-value=2.97*10^−22^) in the four independent studies can be explained by the UKB GWAS results. In conclusion, our study identifies many new genetic variants at SNP, locus and gene levels and advances our understanding of the pleiotropy and genetic co-architecture between brain volumes and other traits.

---

Regional brain volumes are heritable measures of brain functional and structural changes. Volumetric variations of human brain are known to be phenotypically and genetically associated with heritable cognitive and mental health traits (1–5), and it is an active research area to understand the shared genetic influences in these traits (6). Individual variations of human brain volume are usually quantified by magnetic resonance imaging (MRI). In region of interest (ROI)-based analysis, whole brain MRIs are processed and annotated onto many per-defined ROIs, and then regional volumetric phenotypes are generated to measure the structure of brain ROIs. Family and population-based studies have both shown that these volumetric phenotypes are highly heritable (7–9), and common single-nucleotide polymorphism (SNP) markers collected across the genome can account for a large proportion of phenotypic variation (10). A highly polygenic or omnigenic (11, 12) genetic architecture has been observed, which indicates that a large number of genetic variants influence regional brain volumes and their genetic contributions are widespread across the whole genome.

Several genome-wide association studies (GWAS) (3, 7, 8, 13–18) have been conducted to identify genetic risk variants for brain volumetric phenotypes. However, except for the whole brain volume and volumes of few specific ROIs (e.g., hippocampus in subcortical area (3, 8, 19)), GWAS of most brain volumetric phenotypes were insufficiently powered, for which the largest sample size of discovery GWAS was less than 10,000 in (7). Such GWAS sample size is much smaller than those of recent GWAS of other heritable brain-related traits, such as cognitive function (20), neuroticism (21), and intelligence (22), where sample sizes ranged from 269,867 to 449,484. Given the polygenic nature of brain volumes, most of the genetic risk variants may remain undetected, and GWAS with larger sample size can uncover more associated variants and enrich the pleiotropy and genetic co-architecture with other traits. Recently, the UK Biobank (UKB, (23)) study team has collected and released MRI data for more than 20,000 participants. In addition, publicly available imaging genetic datasets also emerge from several other independent studies, including Philadelphia Neurodevelopmental Cohort (PNC, (24)), Alzheimer’s Disease Neuroimaging Initiative (ADNI, (25)), Pediatric Imaging, Neurocognition, and Genetics (PING, (26)), and the Human Connectome Project (HCP,(27)), among others. These datasets provide a new opportunity to perform better-powered GWAS of all ROI brain volumes.

Here we downloaded the raw MRI data from these data resources and processed the data using consistent standard procedures via advanced normalization tools (ANTs, (28)) to generate 101 regional (and total) brain volume phenotypes (referred as ROI volumes), including the total brain volume (TBV), gray matter (GM), white matter (WM), and cerebrospinal fluid (CSF). 19,629 UKB individuals of British ancestry were used in the main discovery GWAS. Other four datasets with relatively small sample sizes (total sample size 2,192 after quality controls) were used to validate the UKB findings and finally a meta-analysis was performed to combine all the data. We started our analysis of UKB data with estimating the SNP heritability, which is the proportion of phenotypic variation that can be explained by the additive effects of all common autosomal SNPs (29). Particularly, the UKB MRI data were released at different time points. We organized them in two parts: the first part was released in 2017 (referred as phase 1 in this paper, n=9,198), most of which has been analyzed in (7), and the second part was released in 2018 (referred as phase 2, n=10,431). To detect any potential heterogeneity of the two phases, we compared the SNP heritability estimated in phase 2 data to those in phase 1 data, which were reported in (10). We then carried out GWAS to identify the associated genetic variants for each ROI volume. We performed gene-based association analysis via MAGMA (30) to uncover gene-level associations, and performed post-GWAS functional mapping and annotation (FUMA, (31)) to explore the functional consequences of the significant SNPs. We calculated the pairwise genetic correlation between ROI volumes and 50 brain-related complex traits by the LD score regression (LDSC, (32)). To confirm the robustness of UKB GWAS findings, we jointly analyzed the UKB GWAS results with those from PNC, ADNI, PING and HCP. We developed genome-wide polygenic risk scores (PRS) to assess the predictive ability of the UKB GWAS results on the other four datasets. GWAS summary statistics of the UKB sample and meta-analysis for the five studies have been made available to public at https://med.sites.unc.edu/bigs2/data/gwas-summary-statistics/.

## RESULTS

### SNP heritability estimates of the two UKB phases data

**Supplementary Fig. 1** compares the SNP heritability (*h^2^*) estimated separately from UKB phase 1 and 2 data. The correlation of these estimates was 0.79, indicating moderate to high level of agreement in terms of the degree of genetic contributions to ROI between the two phases. Six ROIs had >0.6 *h^2^* estimates in both phases, including TBV, cerebellar vermal lobules VIII-X, cerebellar vermal lobules I-V, brain stem, and left/right cerebellum exterior. The *h^2^* estimates from the combined data were highly correlated with those from phase 1 (correlation=0.91) and phase 2 (correlation=0.92) **(Supplementary Figs. 2-3)**. The SNP heritability estimates, standard errors, raw and Bonferroni-corrected p-values from the one-sided likelihood ratio tests are provided in **Supplementary Table 1**. Significant genetic controls widely spread across most ROIs of the whole brain (mean *h^2^*=0.35, *h^2^* range=[0.15,0.71], standard error=0.15). Heritability of left/right basal forebrain (*h^2^*=0.09/0.11) and optic chiasm (*h^2^*=0.01) were insignificant.

### Significant GWAS associations of 101 ROI volumes

We carried out GWAS of the 101 ROI volumes with using 8,944,375 SNPs after genotyping quality controls. Manhattan and QQ plots of all the 101 phenotypes are displayed in **Supplementary Fig. 4**. There were 22,353 significant associations at the conventional 5*10^−8^ GWAS significance level and 12,060 significant ones at the 4.9*10^−10^ significance level (that is, 5*10^−8^/101, additionally adjusted for all 101 GWAS performed) **(Supplementary Fig. 5, Supplementary Table 2)**. TBV had the largest number of significant associations, which was 3,408 at 4.9*10^−10^ significance level. In addition to TBV, left/right hippocampus, left/right putamen, and cerebellar vermal lobules VIII-X had more than 500 significant associations. In the rest of this paper, we refer 4.9*10^−10^ as the significance threshold for SNP-level associations unless otherwise stated.

287 independent significant SNPs had 392 significant associations with 54 ROIs (**Supplementary Table 3**). Independent significant SNPs were defined as significant SNPs that were independent of other significant SNPs by FUMA (Online Methods, (31)). The number of associations for each ROI is displayed in Figure 1 and **Supplementary Table 4.** Left/right hippocampus, cerebellar vermal lobules VIII-X, left/right putamen, and cerebellar vermal lobules I-V had at least 19 independent significant SNPs. Other ROIs that had at least 10 independent significant SNPs included left/right precentral, brain stem, X4th ventricle, left/right lateral ventricle, left/right cerebellum white matter, and TBV. The number of independent significant associations on each chromosome is shown in **Supplementary Table 5**, and clearly chromosome 12 had the largest number of SNP-level associations with ROI volumes **(Supplementary Fig. 6)**.

**Figure 1.**
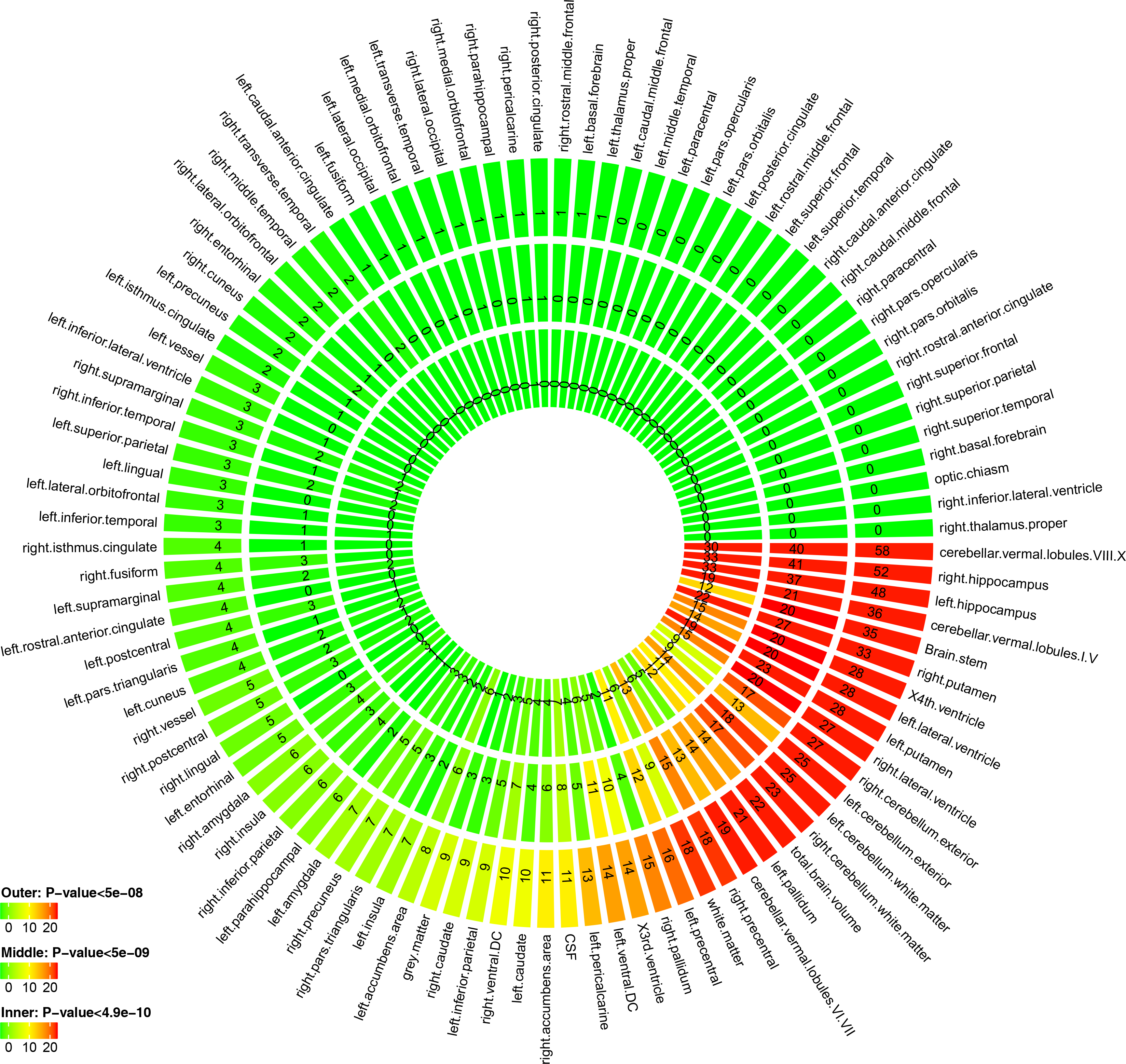
Number of independent significant SNP associations discovered in UKB GWAS at different significance levels. Outer layer: p-value <5*10^−8^; middle layer: p-value <5*10^−9^; and inner layer: p-value <4.9*10^−10^.

The 392 independent significant SNP-level associations can be further characterized (Online Methods) as 134 significant associations between genetic risk loci and ROI volumes (Table 1, **Supplementary Table 6)**. Brain stem, cerebellar vermal lobules VIII-X, left/right lateral ventricle, TBV and WM had at least five genetic risk loci **(Supplementary Table 7)**. Each chromosome had at least one genetic risk locus except for chromosomes 13 and 21 **(Supplementary Tables 8)**. Results at significance thresholds 5*10^−8^ and 5*10^−9^ are also provided in above tables and are summarized in **Supplementary Table 9**.

**Table 1.** List of significant genetic risk loci identified by UKB GWAS at 4.9*10^−10^ significant level. RSID: rsID of the top lead SNP; Position: position of top lead SNP.

| RSID | ROI | CHR | Position | p-value | p-value<br>(second ROI, if<br>any) | p-value<br>(third ROI, if<br>any) | Start position of the locus | End position of the locus |
| --- | --- | --- | --- | --- | --- | --- | --- | --- |
| rs2817145 | left.caudate | 1 | 3133422 | 9.29E-16 |  |  | 3118674 | 3149789 |
| rs2817145 | right.caudate | 1 | 3133422 | 7.69E-13 |  |  | 3121877 | 3149789 |
| rs3120124 | left.pallidum | 1 | 43764165 | 7.95E-11 |  |  | 43760070 | 43949718 |
| rs6658111 | X4th.ventricle | 1 | 47980916 | 2.60E-18 |  |  | 47712057 | 47980916 |
| rs6658111 | left.parahippocampal | 1 | 47980916 | 2.62E-16 |  |  | 47915175 | 47993332 |
| rs6658111 | CSF | 1 | 47980916 | 3.58E-10 |  |  | 47974123 | 47980916 |
| rs3176459 | left.amygdala | 1 | 51437247 | 1.48E-10 |  |  | 50828316 | 51526458 |
| rs12072311 | left.cerebellum.exterior | 1 | 51572465 | 1.51E-12 |  |  | 50882506 | 52215089 |
| rs12072311 | right.cerebellum.exterior | 1 | 51572465 | 5.49E-14 |  |  | 50882506 | 52368501 |
| rs74091739 | cerebellar.vermal.lobules.VIII.X | 1 | 76028726 | 3.21E-45 |  |  | 75818830 | 76095959 |
| rs2748444 | cerebellar.vermal.lobules.I.V | 1 | 76010959 | 1.23E-19 |  |  | 75946751 | 76091842 |
| rs76934732 | cerebellar.vermal.lobules.VI.VII | 1 | 76013268 | 2.72E-11 |  |  | 75946751 | 76073887 |
| rs11392431,<br>rs2991716 | right.inferior.parietal,<br>left.inferior.parietal | 1 | 88423056,<br>88423966 | 3.10E-10 | 3.22E-10 |  | 88196463 | 88435308 |
| rs1044595 | white.matter | 1 | 180943529 | 4.64E-13 |  |  | 180940588 | 181017348 |
| rs1452628 | grey.matter | 1 | 215139887 | 7.34E-12 |  |  | 215134041 | 215327772 |
| rs1030088 | left.inferior.temporal | 2 | 65937745 | 4.22E-10 |  |  | 65937041 | 65999623 |
| rs34111434 | right.transverse.temporal | 2 | 150048607 | 3.60E-10 |  |  | 149945701 | 150084200 |
| rs5835889 | right.hippocampus | 2 | 162845572 | 3.78E-10 |  |  | 162314229 | 162891848 |
| rs5835889 | left.hippocampus | 2 | 162845572 | 9.80E-12 |  |  | 162796517 | 162891848 |
| rs759663 | cerebellar.vermal.lobules.VIII.X | 2 | 208080975 | 2.76E-12 |  |  | 208017033 | 208102505 |
| rs2234675 | left.cerebellum.exterior | 2 | 223085955 | 1.19E-10 |  |  | 223046616 | 223085955 |
| rs12636275 | right.inferior.temporal | 3 | 89523038 | 9.00E-11 |  |  | 89451721 | 89751353 |
| rs12633109 | right.vessel | 3 | 146392141 | 2.89E-11 |  |  | 146337667 | 146396844 |
| rs2279829 | right.pars.triangularis | 3 | 147106319 | 4.52E-14 |  |  | 147090583 | 147224629 |
| rs2279829 | left.pars.triangularis | 3 | 147106319 | 3.56E-10 |  |  | 147095294 | 147224629 |
| rs2279829 | left.postcentral | 3 | 147106319 | 1.87E-10 |  |  | 147095294 | 147140680 |
| rs2279829 | right.postcentral | 3 | 147106319 | 7.02E-12 |  |  | 147095294 | 147268599 |
| rs34859790, rs61500084,<br>rs61500084 | left.inferior.lateral.ventricle,<br>left.lateral.ventricle,<br>right.lateral.ventricle | 3 | 190652830,<br>190636749,19063<br>6749 | 4.39E-11 | 2.80E-13 | 2.59E-17 | 190591418 | 190678743 |
| rs10706986 | left.accumbens.area,<br>right.accumbens.area | 3 | 190646282 | 2.04E-10 | 1.40E-13 |  | 190592836 | 190678743 |
| rs13073466 | cerebellar.vermal.lobules.VIII.X | 3 | 192617930 | 1.38E-10 |  |  | 192613518 | 192716379 |
| rs10034561 | cerebellar.vermal.lobules.VIII.X | 4 | 66701964 | 2.28E-16 |  |  | 66629087 | 66966038 |
| rs17755998 | cerebellar.vermal.lobules.VI.VII | 4 | 66714289 | 1.08E-21 |  |  | 66653473 | 67034895 |
| rs13107325 | left.accumbens.area | 4 | 103188709 | 5.05E-19 |  |  | 102702364 | 103387161 |
| rs13107325 | right.accumbens.area | 4 | 103188709 | 9.64E-16 |  |  | 102702364 | 103388441 |
| rs13107325 | white.matter | 4 | 103188709 | 2.52E-13 |  |  | 103001649 | 103387161 |
| rs10478102 | cerebellar.vermal.lobules.VIII.X | 5 | 112398770 | 4.99E-15 |  |  | 112379434 | 112442272 |
| rs152234 | white.matter | 5 | 179335060 | 2.73E-12 |  |  | 179287576 | 179370127 |
| rs2819861 | left.putamen | 6 | 45424940 | 8.20E-12 |  |  | 45407654 | 45510107 |
| rs35405209 | right.pallidum | 6 | 45428508 | 2.42E-10 |  |  | 45407654 | 45461253 |
| rs12215796 | Brain.stem | 6 | 55715715 | 4.20E-10 |  |  | 55679959 | 55717728 |
| rs9296804 | left.cerebellum.white.matter | 6 | 55717728 | 4.14E-10 |  |  | 55712608 | 55717728 |
| rs9345125 | right.precentral | 6 | 92002653 | 1.14E-10 |  |  | 92002569 | 92010010 |
| rs2153960 | Brain.stem, left.ventral.DC,<br>right.ventral.DC | 6 | 108988184 | 7.69E-12 | 1.05E-11 | 6.32E-12 | 108861264 | 109020032 |
| rs2764264 | total.brain.volume | 6 | 108934461 | 5.16E-13 |  |  | 108861264 | 109019323 |
| rs11759026 | total.brain.volume | 6 | 126792095 | 1.25E-16 |  |  | 126659043 | 127167072 |
| rs62435771 | CSF | 6 | 169470668 | 3.26E-10 |  |  | 169469707 | 169555989 |
| rs368699386 | left.lateral.ventricle | 7 | 2803883 | 1.09E-14 |  |  | 2745766 | 2912928 |
| rs368699386,<br>rs529794364 | X3rd.ventricle,<br>right.lateral.ventricle | 7 | 2803883,<br>2791033 | 3.28E-12 | 3.17E-12 |  | 2752152 | 2912928 |
| rs151057105 | left.ventral.DC, right.ventral.DC | 7 | 54944920 | 7.38E-14 | 5.51E-14 |  | 54909928 | 55015505 |
| rs4006770 | cerebellar.vermal.lobules.VIII.X | 7 | 103361695 | 8.81E-26 |  |  | 103200019 | 103396186 |
| rs12705150 | cerebellar.vermal.lobules.VI.VII | 7 | 103356275 | 2.53E-11 |  |  | 103356062 | 103373126 |
| rs142005327 | white.matter | 7 | 120969969 | 2.21E-11 |  |  | 120954908 | 121040782 |
| rs7778054 | cerebellar.vermal.lobules.I.V | 7 | 156095814 | 1.40E-11 |  |  | 156066471 | 156170722 |
| rs196814 | left.pallidum | 8 | 24716594 | 2.18E-10 |  |  | 24679371 | 24764896 |
| rs13268108 | right.cerebellum.exterior | 8 | 142039586 | 1.30E-10 |  |  | 142033034 | 142042870 |
| rs11789773 | right.postcentral | 9 | 76157130 | 1.10E-10 |  |  | 75849823 | 76193770 |
| rs12344396 | left.cerebellum.white.matter | 9 | 118921327 | 3.75E-16 |  |  | 118917007 | 119098518 |
| rs1405 | Brain.stem | 9 | 118954624 | 7.48E-16 |  |  | 118917007 | 119002278 |
| rs72754248 | left.cerebellum.exterior,<br>right.cerebellum.exterior | 9 | 119061396 | 7.72E-39 | 1.25E-37 |  | 118917007 | 119134875 |
| rs72754266 | right.cerebellum.white.matter | 9 | 119093757 | 2.51E-15 |  |  | 118917007 | 119098518 |
| rs147269950 | X4th.ventricle | 9 | 119098518 | 1.00E-14 |  |  | 119061396 | 119098518 |
| rs7030607 | left.hippocampus,<br>right.hippocampus | 9 | 119245183 | 6.81E-19 | 1.37E-14 |  | 119241165 | 119479868 |
| rs77641763 | cerebellar.vermal.lobules.VIII.X | 9 | 140265782 | 1.20E-10 |  |  | 140241209 | 140278206 |
| rs4748994 | left.caudate | 10 | 25300609 | 3.10E-11 |  |  | 25233290 | 25377328 |
| rs6584542 | total.brain.volume | 10 | 104995495 | 3.69E-14 |  |  | 104965551 | 105176914 |
| rs4752582 | left.cerebellum.exterior | 10 | 123443605 | 1.82E-10 |  |  | 123443605 | 123443605 |
| rs4046908 | white.matter | 11 | 1483117 | 7.69E-11 |  |  | 1472475 | 1632590 |
| rs10770131 | left.lateral.ventricle,<br>right.lateral.ventricle | 11 | 10682248 | 5.99E-12 | 1.89E-11 |  | 10655021 | 10699750 |
| rs1187162 | left.caudate | 11 | 92011126 | 4.02E-11 |  |  | 92001738 | 92036512 |
| rs1187162 | right.caudate | 11 | 92011126 | 1.38E-10 |  |  | 92001738 | 92315907 |
| rs688968 | left.putamen, right.putamen | 11 | 117406739 | 7.36E-13 | 5.10E-11 |  | 117383215 | 117429845 |
| rs7936928 | CSF | 11 | 130279168 | 1.28E-16 |  |  | 130260673 | 130297957 |
| rs11062908 | left.amygdala, right.amygdala | 12 | 4005222 | 1.42E-16 | 1.34E-16 |  | 4004752 | 4013260 |
| rs17178006 | right.hippocampus | 12 | 65718299 | 2.84E-32 |  |  | 65386792 | 66065136 |
| rs17178006 | left.hippocampus | 12 | 65718299 | 9.50E-33 |  |  | 65400292 | 66065136 |
| rs17178006 | right.amygdala | 12 | 65718299 | 3.93E-10 |  |  | 65718299 | 65874956 |
| rs8756 | total.brain.volume | 12 | 66359752 | 4.42E-14 |  |  | 66326943 | 66389968 |
| rs11111094 | left.cerebellum.white.matter, right.cerebellum.white.matter | 12 | 102328806 | 3.03E-31 | 9.74E-28 |  | 102096012 | 102946220 |
| rs11111094 | Brain.stem | 12 | 102328806 | 3.54E-22 |  |  | 102319002 | 102866662 |
| rs35872193 | right.cerebellum.exterior | 12 | 102442428 | 9.61E-12 |  |  | 102394872 | 102962583 |
| rs12146713 | left.lateral.ventricle, right.lateral.ventricle | 12 | 106476805 | 1.86E-11 | 1.88E-13 |  | 106476805 | 106510413 |
| rs10778498 | Brain.stem | 12 | 107063340 | 5.77E-14 |  |  | 106960182 | 107522203 |
| rs146607495 | left.hippocampus | 12 | 117319202 | 1.24E-23 |  |  | 117229539 | 117506632 |
| rs146607495 | right.hippocampus | 12 | 117319202 | 5.04E-25 |  |  | 117234676 | 117528064 |
| rs17126556 | CSF | 14 | 53942788 | 6.98E-12 |  |  | 53917808 | 53996110 |
| rs8014725 | right.putamen | 14 | 56186953 | 2.81E-32 |  |  | 55921561 | 56206214 |
| rs8017172 | left.putamen | 14 | 56199048 | 2.57E-28 |  |  | 55998163 | 56206214 |
| rs8014725 | left.pallidum | 14 | 56186953 | 7.17E-11 |  |  | 56177508 | 56201197 |
| rs76341705 | left.pericalcarine | 14 | 59628679 | 9.97E-21 |  |  | 59449138 | 59916960 |
| rs76341705 | left.cuneus | 14 | 59628679 | 2.90E-12 |  |  | 59585670 | 59669948 |
| rs147148763, rs5809016 | left.precuneus, right.precuneus | 14 | 59631075, 59627434 | 2.87E-10 | 4.22E-12 |  | 59588323 | 59860859 |
| rs5809016, rs73313052 | left.inferior.parietal, right.pericalcarine | 14 | 59627434, 59625997 | 5.71E-11 | 2.56E-10 |  | 59588323 | 59669948 |
| rs2033939 | right.precentral | 15 | 39633904 | 4.46E-63 |  |  | 39355860 | 39670175 |
| rs2033939 | left.precentral | 15 | 39633904 | 5.11E-73 |  |  | 39534404 | 39670175 |
| rs28520337, rs4924345 | left.supramarginal, right.supramarginal | 15 | 39647894, 39639898 | 1.86E-10 | 2.18E-12 |  | 39617295 | 39664000 |
| rs34680120 | left.superior.parietal | 15 | 39664000 | 5.81E-11 |  |  | 39631771 | 39664000 |
| rs4775006 | X4th.ventricle | 15 | 58215727 | 1.01E-20 |  |  | 58188859 | 58385264 |
| rs12921170 | right.lateral.ventricle | 16 | 87227397 | 4.01E-24 |  |  | 87211783 | 87357985 |
| rs4843552 | left.lateral.ventricle | 16 | 87233516 | 2.93E-22 |  |  | 87211963 | 87266245 |
| rs4843550, rs4843560 | X3rd.ventricle, X4th.ventricle | 16 | 87236383, 87225143 | 1.86E-14 | 4.78E-17 |  | 87220694 | 87265438 |
| rs3833159 | right.precuneus | 17 | 27953390 | 3.89E-11 |  |  | 27925812 | 28553639 |
| rs3110494 | left.insula | 17 | 27976597 | 1.01E-13 |  |  | 27935546 | 28553639 |
| rs3110494 | right.insula | 17 | 27976597 | 5.44E-13 |  |  | 27935546 | 28538715 |
| rs78777685 | cerebellar.vermal.lobules.I.V | 17 | 35257232 | 5.46E-23 |  |  | 35215422 | 35266724 |
| rs118087478 | total.brain.volume | 17 | 44051589 | 1.03E-13 |  |  | 43463493 | 44865603 |
| rs11665242, rs6508230 | left.putamen, right.putamen | 18 | 50907127, 50884924 | 2.25E-15 | 2.52E-15 |  | 50555225 | 51061399 |
| rs60575064 | X3rd.ventricle | 19 | 33565344 | 9.33E-12 |  |  | 33525467 | 33631912 |
| rs35255138 | cerebellar.vermal.lobules.I.V | 19 | 41197268 | 3.51E-11 |  |  | 41148797 | 41197268 |
| rs6121038 | right.pallidum | 20 | 30254773 | 2.20E-12 |  |  | 29420066 | 30437522 |
| rs6121038 | left.pallidum | 20 | 30254773 | 1.17E-12 |  |  | 29509439 | 30439298 |
| rs6121038 | right.putamen | 20 | 30254773 | 3.01E-10 |  |  | 30235302 | 30434084 |
| rs6062237, rs6062264 | right.ventral.DC, left.ventral.DC | 20 | 61154437, 61154871 | 2.55E-12 | 2.16E-14 |  | 61141981 | 61158050 |
| rs34134374 | left.lateral.ventricle | 22 | 38138284 | 4.52E-10 |  |  | 38102776 | 38304858 |

### Concordance with previous GWAS results

We performed association lookups for the 287 independent significant SNPs and their correlated SNPs in genetic risk loci (Online Methods) on the NHGRI-EBI GWAS catalog (33). We found that 117 independent significant SNPs (associated with 36 ROI volumes) have previously reported GWAS associations with any traits **(Supplementary Table 10)**. Our results tagged many SNPs that were previously reported in GWAS of ROI volumes, including 14 SNPs in van der Meer, Rokicki (3) for hippocampal subfield volumes, 11 in Hibar, Stein (8) for subcortical brain region volumes, 5 in Chen, Wang (34) for putamen volume, 4 in Bis, DeCarli (18) for hippocampal volume, 2 in Hibar, Adams (14) for hippocampal volume, 2 in Stein, Medland (35) for brain structure, 2 in Ikram, Fornage (17) for intracranial volume, 1 in Furney, Simmons (36) for whole brain volume, and 1 in Baranzini, Wang (37) for normalized brain volume **(Supplementary Table 11)**. For the other traits, we highlighted previous associations of 29 different SNPs with mental health disease/disorders (such as schizophrenia, autism spectrum disorder [ASD], and depression), 78 with cognitive functions, 17 with educational attainment, 20 with neuroticism, 14 with Parkinson’s disease, 3 with reaction time, and 1 with Alzheimer’s disease. More previous GWAS results were found when the significance threshold was relaxed to 5*10^−8^ **(Supplementary Table 12).** We also compared our results with those reported in (7). Elliott, Sharp (7) performed GWAS of 3,144 imaging phenotypes (including brain volume phenotypes processed by FreeSurfer (38)) using the UKB phase 1 data (n=8,428). When both being corrected for the number of performed GWAS, 26 of the 78 unique SNPs (covered 66 of the 368 significant associations) reported in (7) were within LD of our independent significant SNPs **(Supplementary Table 13)**. When both being relaxed to the 5*10^−8^ significance threshold, 119 of their 616 unique SNPs (covered 493 of the 1,262 significant associations) were within LD of our independent significant SNPs.

### Gene-based association analysis and functional mapping

We performed gene-based association analysis with GWAS summary statistics for 18,796 candidate genes (Online Methods). We found 237 significant gene-level associations (p-value<2*10^−8^, adjusted for multiple traits) between 142 genes and 47 ROIs (Table 2, **Supplementary Table 14).** Our results replicated 29 genes discovered in previous studies, including *FOXO3* in Baranzini, Wang (37), *HMGA2* and *HRK* in Stein, Medland (35), *KANSL1, MAPT, STH* and *CENPW* in Ikram, Fornage (17), *SLC44A5* in Furney, Simmons (36), *MSRB3, BCL2L1, DCC, CRHR1* in Hibar, Stein (8), *LEMD3, WIF1* and *ASTN2* in Bis, DeCarli (18), *FAM53B, METTL10* and *FAF1* in van der Meer, Rokicki (3), *DSCAML1* in Chen, Wang (34), *SLC39A1, GATAD2B, DENND4B* in Hibar, Stein (39), and *ZIC4, VCAN, PAPPA, DRAM1, GNPTAB, DAAM1*, and *ALDH1A2* in Elliott, Sharp (7). 13 genes were novel and were not linked to ROI volumes. 57 genes have previously been implicated with cognitive functions, intelligence, education, neuroticism, neuropsychiatric and neurodegenerative diseases/disorders, such as *IGF2BP1* (22, 40, 41), *WNT3* (20, 21, 42, 43), *PLEKHM1* (43–45), and *AGBL2* (21, 43, 46, 47). Particularly, 40 of the 57 pleiotropic genes were novel genes of ROI volumes, and thus these findings substantially uncovered the gene-level pleiotropy between ROI volumes and these traits (Figure 2).

**Figure 2.**
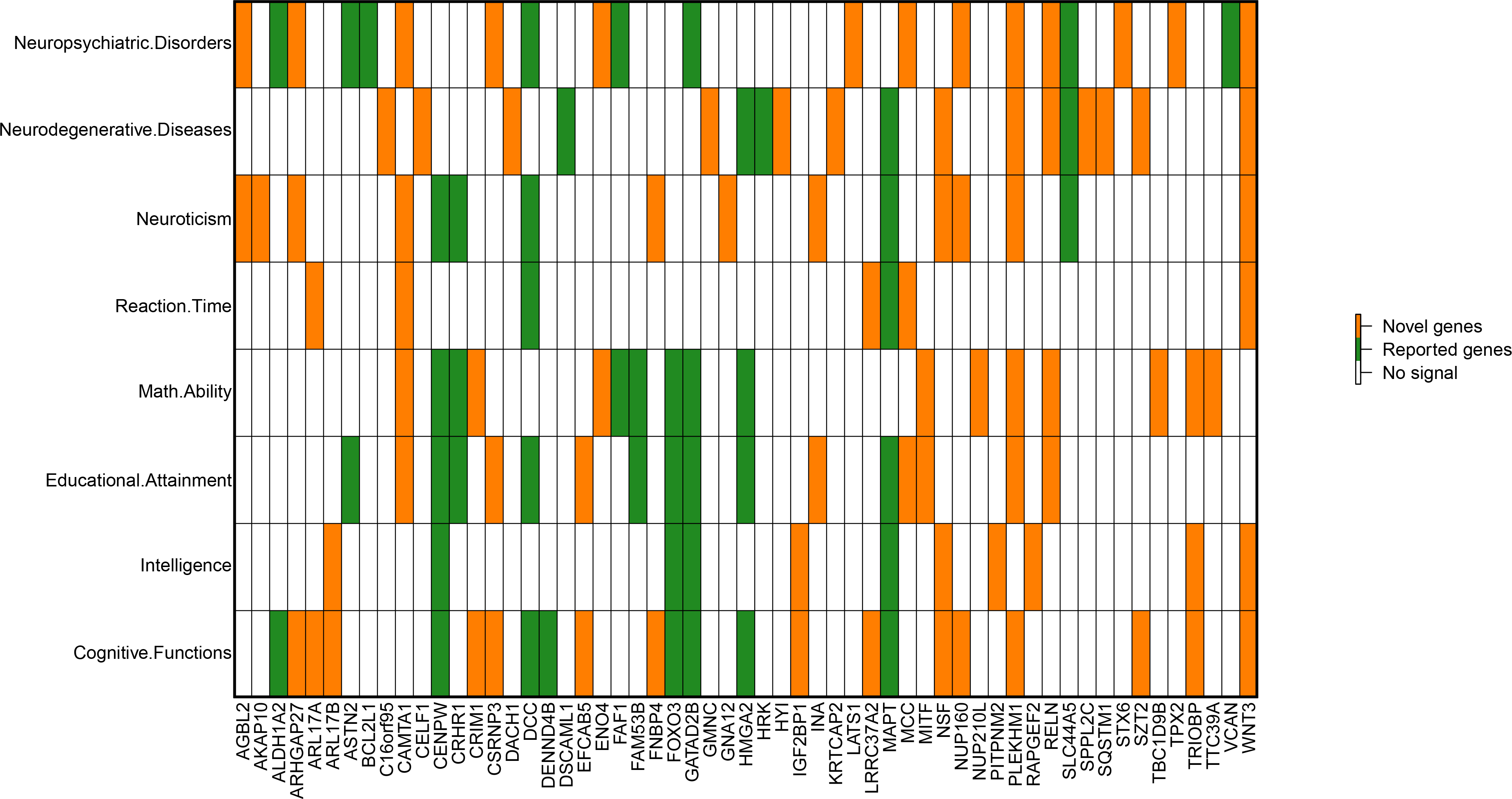
Genes identified in gene-based association analysis of ROI volumes that have been linked to cognitive traits and mental health disease/disorders in previous GWAS.

**Table 2.**
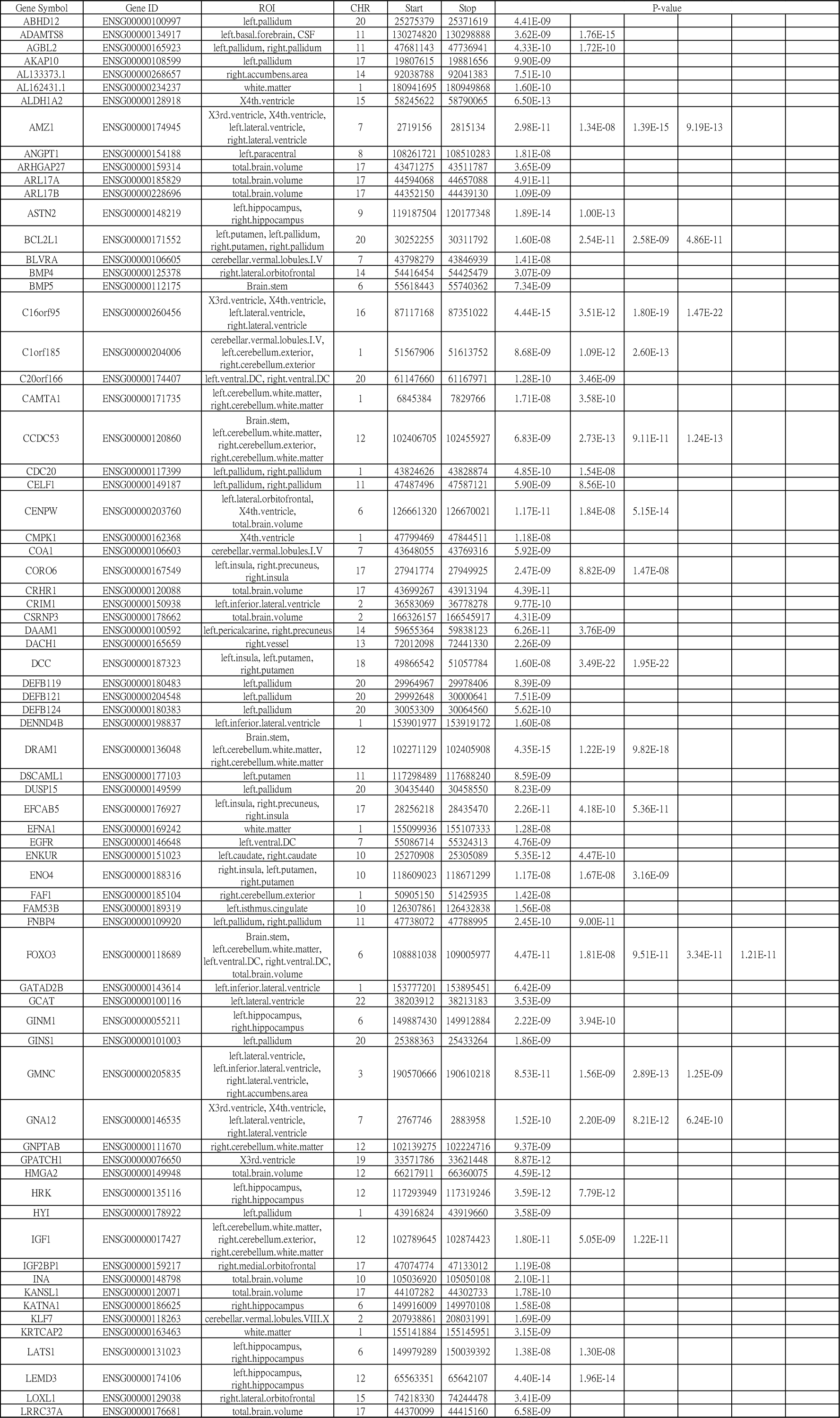
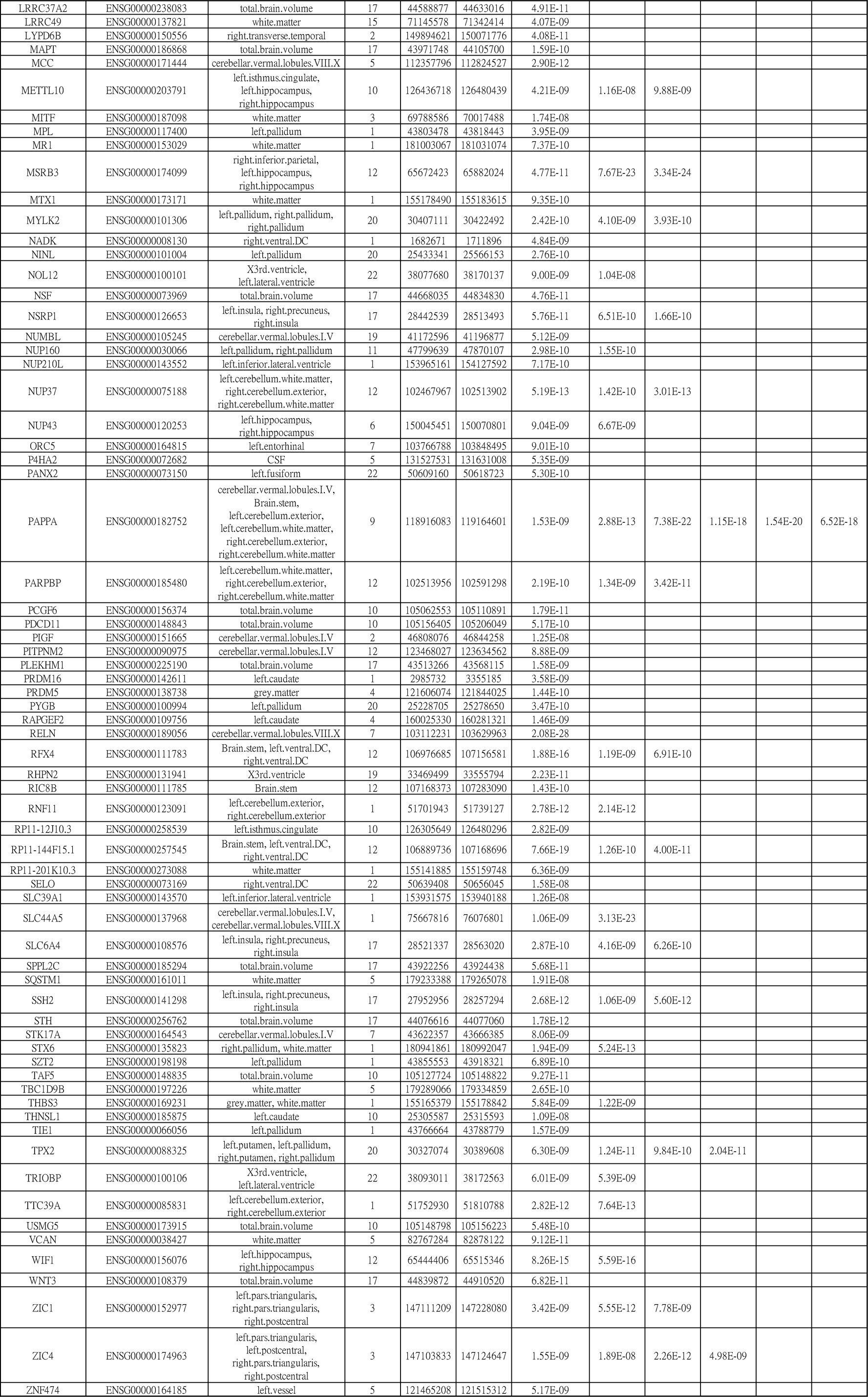
List of significant genes identified by gene-based association analysis in UKB data at 2*10^−8^ significant level.

The independent significant SNPs were also annotated by functional consequences on gene functions **(Supplementary Table 15, Supplementary Fig. 7)**, and were subsequently mapped to genes according to physical position, expression quantitative trait loci (eQTL) association (for brain tissues), and 3D chromatin (Hi-C) interaction (Online Methods). Functional gene mapping yielded 389 significant associations on 214 genes and 49 ROIs **(Supplementary Table 16)**. 122 genes were not discovered in the above gene-based association analysis, which replicated more previous findings on ROI volumes, such as *TBPL2* and *KTN1* in Chen, Wang (34), *FAT3* in Hibar, Stein (8), *SLC4A10, RNFT2, TESC, DMRTA2, CDKN2C* and *DPP4* in van der Meer, Rokicki (3), and *EPHA3, SLC39A8, BANK1, WNT16, CHPT1, ACADM, FAM3C, FBXW8, L3HYPDH, JKAMP*, and *AQP9* in Elliott, Sharp (7). 31 (23 new) of the 122 genes were associated with cognitive functions, intelligence, education, neuroticism, neuropsychiatric and neurodegenerative disorders, such as *NT5C2* (21, 44, 48, 49), *ADAM10* (49, 50), and *GOSR1* (20, 44) **(Supplementary Fig. 8)**.

Gene-priority analysis was performed for 14 brain tissues to examine whether the tissue-specific gene expression levels were related to the associations between genes and ROI volumes (Online Methods). After adjusting for multiple testing (that is, 14*101=1,414 tests) by the Benjamini-Hochberg (B-H) procedure (51) at 0.05 level, we detected nine significant associations, including gene expression in brain hippocampus tissue and gene’s association significance with left hippocampus volume, and gene expression in brain cerebellar hemisphere and cerebellum tissues and gene’s association significance with pallidum and putamen volumes (p-value<2.46*10^−4^) **(Supplementary Table 17)**. These results showed that genes with higher transcription levels on these brain tissues also had stronger associations with the corresponding brain ROI volumes.

### Joint analysis with four independent datasets

To validate the UKB GWAS results, we repeated GWAS of 101 ROI volumes separately on data obtained from four other independent studies: PNC (n=537), HCP (n=334), PING (n=461), and ADNI (n=860). Due to the small sample size of these four datasets, the probability of replicating significant findings in the UKB was low. Instead, we checked whether the SNP effect signs were concordant in the five studies and whether the p-value of top UKB SNPs decreased after meta-analysis (Online Methods). Smaller p-values after meta-analysis indicates similar SNP effects in independent samples (52, 53).

The joint analysis was carried out on 3,841,911 SNPs which were present in all five sets of GWAS results. For the 5,940 significant associations (at 4.9*10^−10^ significance level), 64.6% (3,839) associations had the same effect signs across the five studies, and 97.5% (5,791) associations had the same effect signs in at least four studies (including UKB). 94.0% (1,880) of the top 2,000 significant associations had smaller p-value after meta-analysis, and 92.3% (5,484) of all the 5,940 associations were enhanced. We then performed meta-analysis on all the 8,944,375 UKB GWAS SNPs (SNPs were allowed to be missing in the four independent datasets). There were more significant associations after meta-analysis: 25,083 significant associations at 5*10^−8^ significance level and 14,004 at 4.9*10^−10^ significance level **(Supplementary Table 18, Supplementary Fig. 9)**.

### Genetic correction with other traits

The meta-analysis GWAS results were used to estimate the genetic correlation (gc) with other traits via LDSC (32). As positive controls, we first estimated the genetic correlation between several UKB ROIs volumes (TBV, left/right thalamus proper, left/right caudate, left/right putamen, left/right pallidum, left/right hippocampus, left/right accumbens area) and their corresponding traits studied in the ENIGMA consortium (54). The gc estimates were all significant (p-value<1.20*10^−5^) and average correlation was 0.93 **(Supplementary Table 19).** We then collected 50 sets of publicly available GWAS summary statistics **(Supplementary Table 20)** and calculated their pairwise genetic correlation with ROI volumes **(Supplementary Tables 21)**. We mainly focused on traits that showed evidence of pleiotropy in association lookups. There were 29 significant associations after adjusting for multiple testing (4,900 tests) by the B-H procedure at 0.05 level **(Supplementary Tables 22, Supplementary Fig. 10)**.

Significant genetic correlations linked 16 ROI volumes with general cognitive functions, education (education years, college completion), intelligence, numerical reasoning, reaction time, depressive symptoms, neuroticism, worry, ASD, and bipolar disorder (BD) (Figure 3), which matched our findings in SNP and gene level lookups. Particularly, TBV had positive correlations with cognitive functions, education, intelligence, and numerical reasoning (gc range=[0.20,0.24], mean=0.22, p-value range=[1.54*10^−11^,2.73*10^−5^]). Left posterior cingulate showed positive correlations with cognitive functions, intelligence, and numerical reasoning (gc range=[0.16,0.17], p-value range=[6.09*10^−5^,1.85*10^−4^]). We note that TBV has been adjusted in GWAS of ROIs other than TBV. Right rostral anterior cingulate showed positive correlation with ASD (gc=0.32, p-value=2.00*10^−4^), left rostral middle frontal had positive correlation with BD (gc=0.20, p-value=1.00*10^−4^), and right precuneus had positive correlation with neuroticism (gc=0.17, p-value=1.20*10^−4^). Reaction time had positive genetic correlations with left/right lateral ventricle and X3rd ventricle (gc range=[0.16,0.18], p-value range=[3.13*10^−5^,1.58*10^−4^]), and had negative correlations with left/right pallidum, left/right ventral DC, and white matter (gc range=[−0.20, −0.15], p-value range=[3.64*10^−7^,1.44*10^−5^]). Negative genetic correlations were also found on depressive symptoms (gc=-0.25, p-value=3.33*10^−5^), neuroticism (gc=-0.14, p-value=2.20*10^−4^), and worry (gc=-0.14, p-value=2.94*10^−4^).

**Figure 3.**
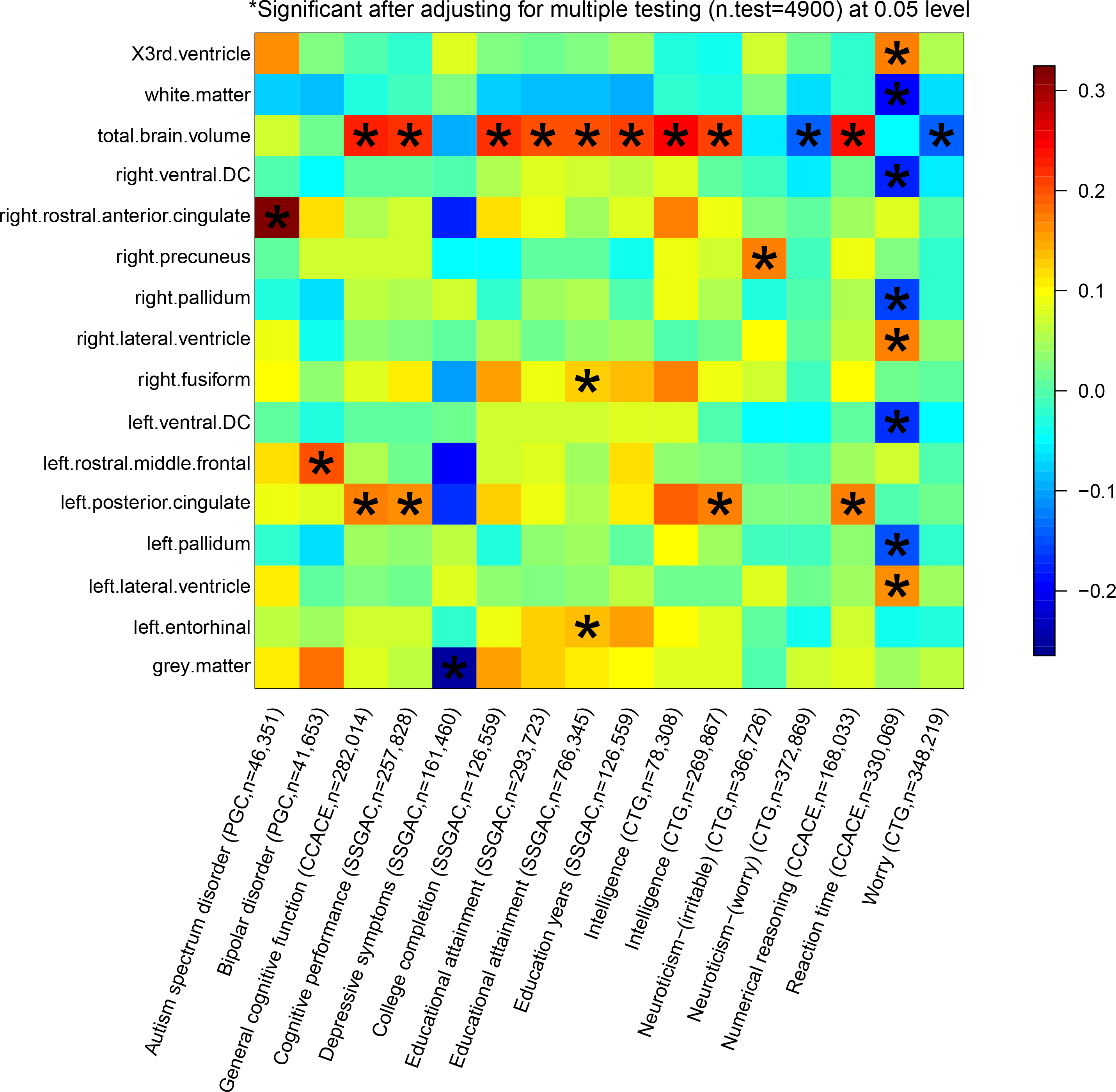
Selected pairwise genetic correlations between ROI volumes and other traits

### Predictive ability of the UKB GWAS results

We examined the out-of-sample prediction power of the UKB GWAS summary statistics using polygenic risk scores prediction (55). We focused the analysis on total brain volume. We first used a ten-fold cross-validation design to examine the prediction power within the UKB sample (Online Methods). Five polygenic profiles were created with p-value thresholds 1, 0.5, 0.05, 5*10^−4^ and 5*10^−8^, respectively, and we examined the incremental R-squared (Online Methods). The PRS can explain 1.51% of the variance in total brain volume (p-value=4.42*10^−110^) **(Supplementary Table 23)**. We then used the GWAS summary statistics of 19,629 UKB individuals to construct polygenic profiles on subjects in PNC, HCP, PING, and ADNI. The UKB-derived PRS were all significantly associated with the phenotype in all the four independent datasets, and can account for 1.38%-4.36% phenotypic variation (p-value range=[2.97*10^−22^, 1.44*10^−6^]). The largest R-squared 4.36% was in PNC dataset with threshold 1 and 224,657 SNP predictors.

## DISCUSSION

In this study, we presented GWAS of 101 ROI volumes using data of 19,629 UKB individuals. Our novel contributions include 1) identification of many newly associated genetic variants at SNP, locus, and gene levels; 2) revealing the genetic co-architecture of brain volume phenotypes and other brain-related complex traits; 3) validation of the UKB results in independent studies; and 4) assessment of the predictive power of UKB GWAS results. Significant (p-value<4.9*10^−10^) associations were found for 54 of the 101 ROIs. With larger sample size, the present study replicated many known genetic variants but also prioritized new ones. Compared to (7), our GWAS not only discovered more genetic variants, but also enriched the degree of (statistical) pleiotropy (56) of the associated genes and characterized the shared genetic influences with cognitive and mental health traits.

However, the current GWAS sample size of ROI volumes (and many other brain imaging phenotypes) is still far from being sufficient. The highly polygenic genetic architecture of ROI volumes requires a larger number of subjects to identify many weak causal SNPs. In the era of sharing GWAS summary statistics, well powered GWAS is essential for ROI volumes to be linked to the genetic co-architecture atlas with other complex traits. For example, a recent study of Watanabe, Stringer (56) to discover the global overview of genetic co-architecture of 2,965 traits only focused on GWAS with sample size larger than 50,000, with the average sample size of selected traits being 256,276. In our genetic correlation analysis, we only obtained limited number of significant correlations, even though many pleiotropic genes were found in association lookups. Therefore, we expect that GWAS of ROI volumes with larger sample size will be available and can further improve our understating of genetic overlaps underlying other traits. Besides increasing the sample size, combining SNP data with external omic information, such as gene expression data (57), may also help elucidate the causal mechanism, improve the prediction performance of SNP data and reveal the genetic connections among traits.

### URLs

ANTs, http://stnava.github.io/ANTs/;

PLINK, https://www.cog-genomics.org/plink2/;

GCTA, http://cnsgenomics.com/software/gcta/;

METAL, https://genome.sph.umich.edu/wiki/METAL;

FUMA, http://fuma.ctglab.nl/;

MGAMA, https://ctg.cncr.nl/software/magma;

LD Score Regression, https://github.com/bulik/ldsc/;

LD Hub, http://ldsc.broadinstitute.org/ldhub/;

MaCH-Admix, http://www.unc.edu/~yunmli/MaCH-Admix;

NHGRI-EBI GWAS Catalog, https://www.ebi.ac.uk/gwas/home;

The atlas of GWAS Summary Statistics, http://atlas.ctglab.nl/;

UK Biobank, http://www.ukbiobank.ac.uk/resources/;

PING, http://pingstudy.ucsd.edu/resources/genomics-core.html;

PNC, https://www.ncbi.nlm.nih.gov/projects/gap/cgi-bin/study.cgi?study_id=phs000607.v1.p1;

ADNI, http://adni.loni.usc.edu/data-samples/;

HCP, https://www.humanconnectome.org/.

## METHODS

Methods are available in the ***Online Methods*** section.

*Note: One supplementary information pdf file and one supplementary zip file are available.*

## ACKNOWLEDGEMENTS

This research was partially supported by U.S. NIH grants MH086633 and MH116527, and a grant from the Cancer Prevention Research Institute of Texas. We thank the individuals represented in the UK Biobank, ADNI, HCP, PING and PNC datasets for their participation and the research teams for their work in collecting, processing and disseminating these datasets for analysis. This research has been conducted using the UK Biobank resource (application number 22783), subject to a data transfer agreement. We gratefully acknowledge all the studies and databases that made GWAS summary data available. Part of data collection and sharing for this project was funded by the Alzheimer’s Disease Neuroimaging initiative (ADNI) (National Institutes of Health Grant U01 AG024904) and DOD ADNI (Department of Defense award number W81XWH-12-2-0012). ADNI is funded by the National Institute on Aging, the National Institute of Biomedical Imaging and Bioengineering and through generous contributions from the following: Alzheimer’s Association; Alzheimer’s Drug Discovery Foundation; Araclon Biotech; BioClinica, Inc.; Biogen Idec Inc.; Bristol-Myers Squibb Company; Eisai Inc.; Elan Pharmaceuticals, Inc.; Eli Lilly and Company; EuroImmun; F. Hoffmann-La Roche Ltd and its affiliated company Genentech, Inc.; Fujirebio; GE Healthcare; IXICO Ltd; Janssen Alzheimer Immunotherapy Research & Development, LLC; Johnson & Johnson Pharmaceutical Research & Development LLC; Medpace, Inc.; Merck & Co., Inc.; Meso Scale Diagnostics, LLC; NeuroRx Research; Neurotrack Technologies; Novartis Pharmaceuticals Corporation; Pfizer Inc.; Piramal Imaging; Servier; Synarc Inc.; and Takeda Pharmaceutical Company. The Canadian Institutes of Health Research is providing funds to support ADNI clinical sites in Canada. Private sector contributions are facilitated by the Foundation for the National Institutes of Health (www.fnih.org). The grantee organization is the Northern California Institute for Research and Education, and the study is coordinated by the Alzheimer’s Disease Cooperative Study at the University of California, San Diego. ADNI data are disseminated by the Laboratory for Neuro Imaging at the University of Southern California. Part of the data collection and sharing for this project was funded by the Pediatric Imaging, Neurocognition and Genetics Study (PING) (U.S. National Institutes of Health Grant RC2DA029475). PING is funded by the National Institute on Drug Abuse and the Eunice Kennedy Shriver National Institute of Child Health & Human Development. PING data are disseminated by the PING Coordinating Center at the Center for Human Development, University of California, San Diego. Support for the collection of the PNC datasets was provided by grant RC2MH089983 awarded to Raquel Gur and RC2MH089924 awarded to Hakon Hakonarson. All PNC subjects were recruited through the Center for Applied Genomics at The Children’s Hospital in Philadelphia. HCP data were provided by the Human Connectome Project, WU-Minn Consortium (Principal Investigators: David Van Essen and Kamil Ugurbil; 1U54MH091657) funded by the 16 NIH Institutes and Centers that support the NIH Blueprint for Neuroscience Research; and by the McDonnell Center for Systems Neuroscience at Washington University.

## AUTHOR CONTRIBUTIONS

B.Z., H.Z., and Y.L. designed the study. B.Z. and T.L. performed the experiments and analyzed the data. T.L., J.Z. Y.S., X.W., L.Y., F.Z., and Z.Z. downloaded the datasets, preprocessed MRI and SNP data, and undertook the quantity controls. B.Z., H.Z., and Y.L. wrote the manuscript with feedback from all authors.

## COMPETETING FINANCIAL INTERESTS

The authors declare no competing financial interests.

## ONLINE METHODS

### GWAS participants and phenotypes

We performed GWAS separately on five publicly available datasets: the UK Biobank (UKB) study, the Human Connectome Project (HCP) study, the Pediatric Imaging, Neurocognition, and Genetics (PING) study, the Philadelphia Neurodevelopmental Cohort (PNC) study, and the Alzheimer’s Disease Neuroimaging Initiative (ADNI) study. The main GWAS made use of data of 19,629 individuals of British ancestry from the UKB study, and the other four GWAS were performed on individuals of European ancestry, see **Supplementary Table 24** for a summary of sample size of each GWAS.

The raw MRI, covariates and SNP data were downloaded from each data resource. We processed the MRI data locally using consistent procedures via advanced normalization tools (ANTs) to generate ROI volume phenotypes for each dataset. The processing steps were detailed in Zhao, Ibrahim (10) and we removed three ROIs (X5th ventricle and left/right lesion) with missing rates > 99%. For each phenotype and continuous covariate variable, we further removed values greater than five times the median absolute deviation from the median value. All individuals were aged between 3 and 92 years. More information about study cohorts can be found in **Supplementary Table 25** and **Supplementary Note**.

### Heritability estimation and Genome-wide association analysis

We estimated the proportion of variation explained by all autosomal SNPs in UKB with using univariate GCTA-GREML analysis (40). The adjusted covariates included age (at imaging), age-squared, gender, age-gender interaction, age-squared-gender interaction, TBV (for ROIs other than TBV itself), as well as the top 40 genetic principle components (PCs) provided by UKB ((58), Data-Field 22009). We performed GWAS for each ROI volume with PLINK (59). The same set of covariates as in GCTA-GREML analysis were adjusted. GWAS were also separately performed on PING, PNC, ADNI, and HCP data. In these four datasets, we adjusted for age, age-squared, gender, age-gender interaction, age-squared-gender interaction, TBV (for ROIs other than TBV itself), and top ten genetic PCs estimated from the SNP data. We also adjusted for the Alzheimer’s disease status in ADNI GWAS.

### Genomic risk loci characterization and comparison with previous findings

Genomic risk loci were defined using FUMA (31) online platform (version: 1.3.4). We input the UKB GWAS summary statistics obtained from PLINK (59). FUMA first identified independent significant SNPs, which were defined as SNPs with a p-value smaller than the predefined threshold and independent of other significant SNPs at R-squared < 0.6. Using these independent significant SNPs, FUMA then constructed LD block for independent significant SNPs by tagged all SNPs that had a MAF ≥ 0.0005 and were in LD (R-squared≥0.6) with at least one of the independent significant SNPs. These SNPs included those from the 1000 Genomes reference panel and may not have been included in the present study. Based on these independent significant SNPs, (independent) lead SNPs were also identified as those that were independent from each other (R-squared<0.1). If LD blocks of independent significant SNPs were closed (<250 kb based on the closest boundary SNPs of LD blocks), they were merged to a single genomic locus. Thus, each genomic locus could contain more than one independent significant SNPs and lead SNPs. More details can be found in Watanabe, Taskesen (31). Independent significant SNPs and all the tagged SNPs were subsequently searched by FUMA on NHGRI-EBI GWAS catalog ((33), version: 2019-01-31) to look for their reported SNP associations (p-value<9*10^−6^) with any traits.

### Gene-based association analysis, functional annotation and gene-property analysis

Gene-based association analysis was carried out for 18,796 protein-coding genes using MAGMA (30), which was also implemented in FUMA (31). SNPs were mapped according to their psychical positions, then the gene-based p-values were calculated by the GWAS summary statistics of mapped SNPs.

In functional annotation and mapping analysis, SNPs signals were annotated with their biological functionality and then were linked to genes by a combination of positional, eQTL, and 3D chromatin interaction mappings (31). Specifically, independent significant SNPs and all the tagged SNPs were first annotated for functional consequences on gene functions (e.g., intergenic, intronic, exonic) using ANNOVAR (60). Functionally-annotated SNPs were then mapped to 35,808 candidate genes based on physical position on the genome (tissue/cell types for 15-core chromatin state: brain), eQTL associations (tissue types: GTEx v7 brain (61), BRAINEAC (62), and CommonMind Consortium (63)) and chromatin interaction mapping (built-in chromatin interaction data: dorsolateral prefrontal cortex, hippocampus (64); annotate enhancer/promoter regions: E053-E082 brain (65)). We used default values for all other parameters.

For the detected genes, we performed lookups on NHGRI-EBI GWAS catalog ((33), version: 2019-01-31) again to explore the previously reported associations with the same or other traits. We focused on traits including cognitive functions (such as general cognitive ability, cognitive performance, and empathy quotient), intelligence, educational attainment, math ability (such as highest math class taken and self-reported math ability), reaction time, neuroticism, neurodegenerative diseases (such as Alzheimer’s disease and Parkinson’s disease), and neuropsychiatric disorders (such as major depressive disorder, Schizophrenia, and bipolar disorder). For the 14 brain tissues (GTEx v7, (61)), we also performed gene-priority analysis via MAGMA (30). That is, for each candidate gene, whether its tissue-specific expression levels can be linked to the strength of its association with ROI volumes.

### Meta-analysis of GWAS results

We meta-analyzed the UKB, PING, PNC, ADNI, and HCP GWAS summary results by METAL (https://genome.sph.umich.edu/wiki/METAL) with the sample-size weighted approach. Since the sample sizes of other four datasets were small, we removed the SNPs that were not presented in the UKB data.

### Genetic correlation estimation with LDSC

The LD Hub (v1.9.1, http://ldsc.broadinstitute.org/ldhub/) was used to estimate the genetic correlation between several UKB ROIs volumes and their corresponding traits studied in the ENIGMA consortium (54). Then the LDSC software (v1.0.0, https://github.com/bulik/ldsc) was used to estimate the pairwise genetic correlation with 50 sets of collected GWAS summary statistics. We used the pre-calculated LD scores provided by LDSC (https://data.broadinstitute.org/alkesgroup/LDSCORE/), which were computed using 1000 Genomes European data. We used HapMap3 SNPs and removed all SNPs on chromosome 6 in the MHC region.

### Polygenic scoring

Polygenic profiles were created to examine the out-of-sample prediction power of the GWAS results. Specifically, we used PLINK (59) to generate risk scores in testing data by summarizing across pruned (window size 50, step 5, R-squared = 0.2) SNP alleles, weighed by their effect sizes estimated from training data. We randomly divided the 19,629 UKB individuals into ten folds, then used nine of these folds as training data to rerun GWAS, and created polygenic profiles on the individuals in the remaining fold, which served as testing data. We repeated this procedure ten times such that each fold alternated to serve as the testing data for exactly one time. Then we used the UKB GWAS results to perform prediction on ADNI, PING, PNC and HCP data. The prediction accuracy was evaluated on all samples in the four testing sets (with phenotype and SNP data available), not limited to individuals of European ancestry used in GWAS. We tried five p-value thresholds for SNP predictor selection: 1, 0.5, 0.05, 5*10^−4^ and 5*10^−8^. The association between polygenic profile and brain volume was estimated and tested in linear regression model, adjusting for the effects of age and gender. The additional variance of brain volume that can be explained by polygenic profile was used to measure the prediction power.

### Data availability

All UKB and meta-analysis GWAS summary statistics of 101 ROI volumes can be found at: https://med.sites.unc.edu/bigs2/data/gwas-summary-statistics/.

