## Supplementary_information for "GWAS of 19,629 individuals identifies novel genetic variants for regional brain volumes and refines their genetic co-architecture with cognitive and mental health traits"

March 21, 2019

#### 1 Supplementary figures

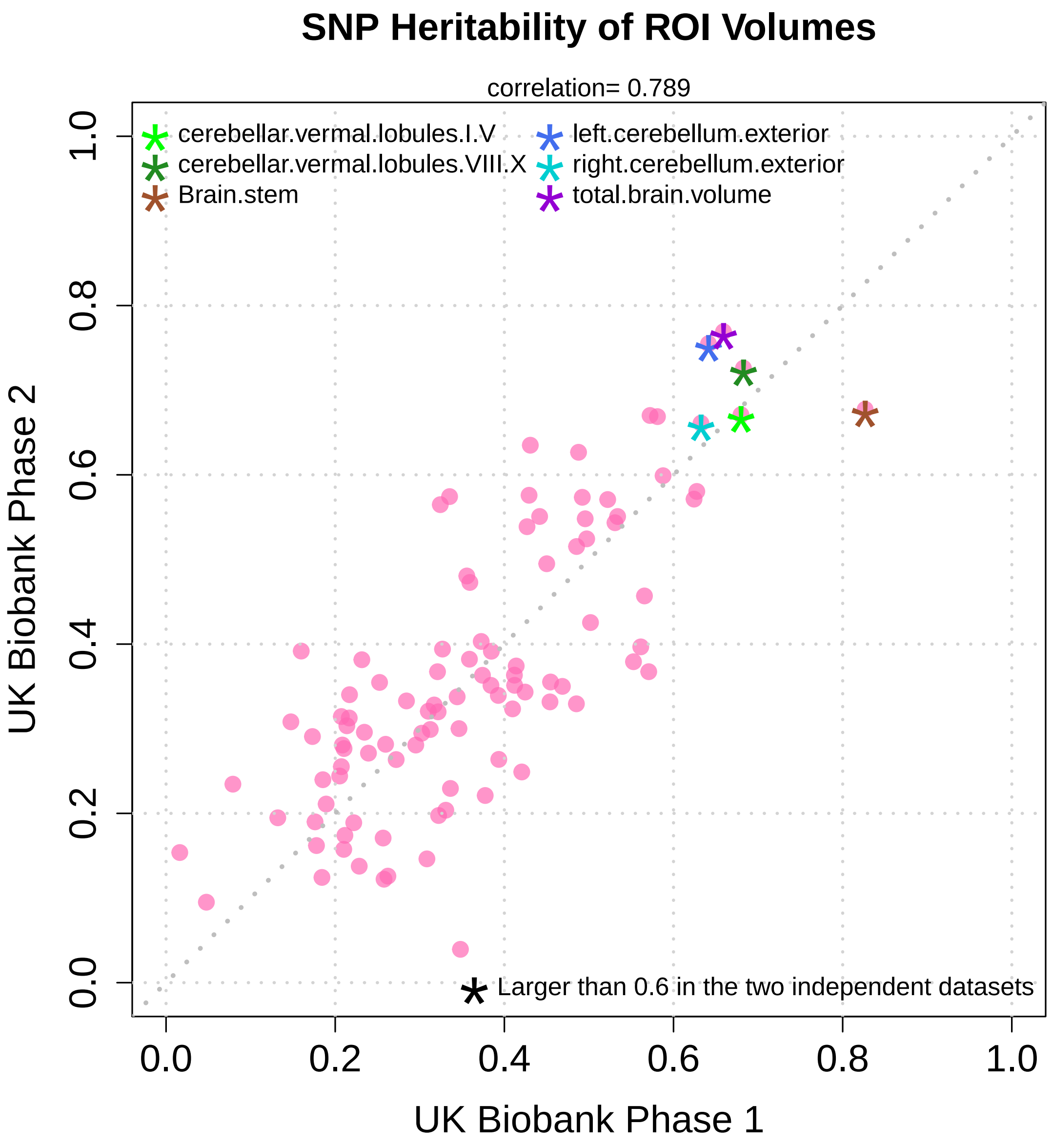


**Supplementary Figure 1:** Comparing SNP heritability estimates of ROI volumes in UK Biobank phase 1 data (n=9,198) and phase 2 data (n=10,431).

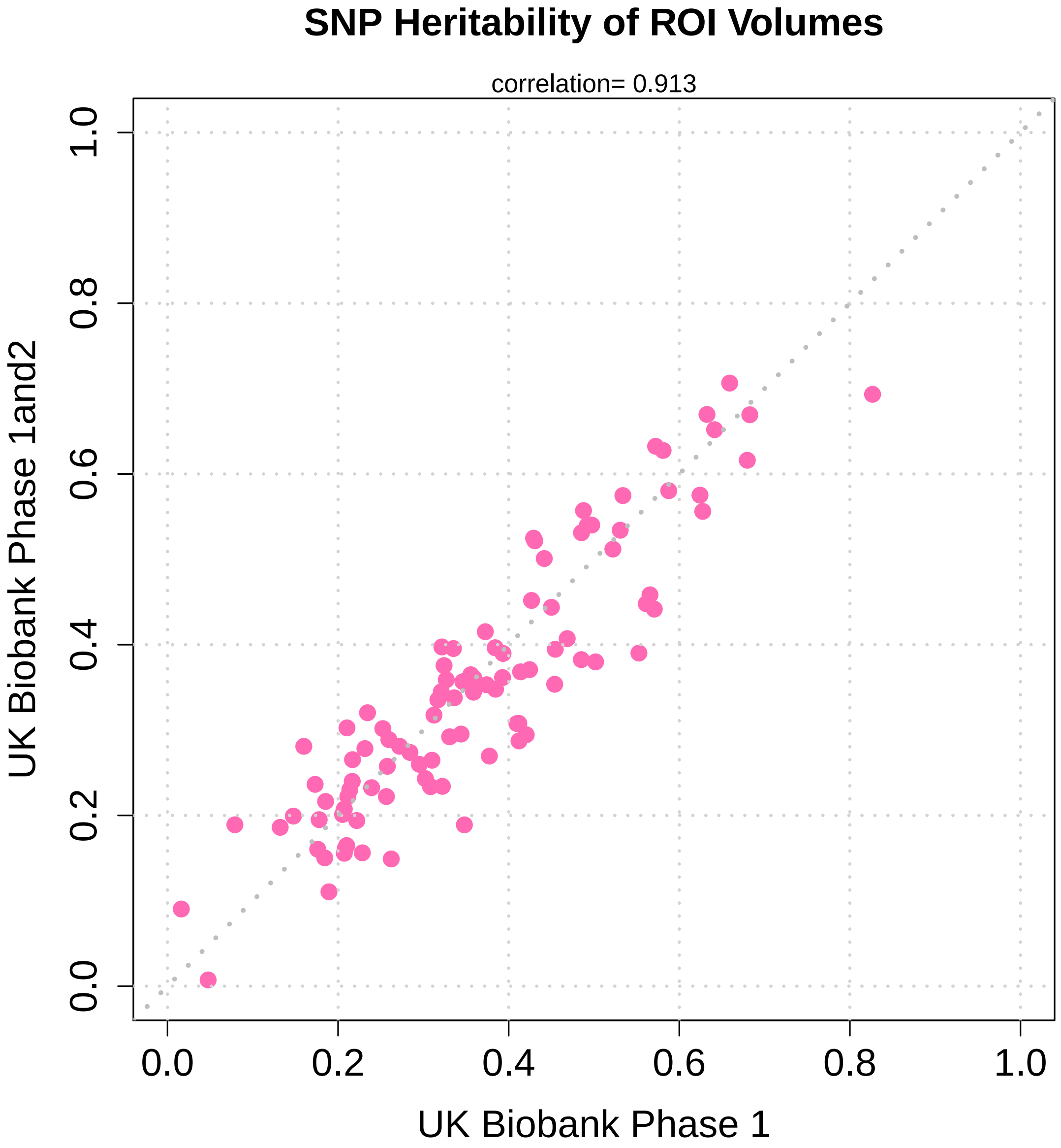


**Supplementary Figure 2:** Comparing SNP heritability estimates of ROI volumes in UK Biobank phase 1 data (n=9,198) and the combined phases 1 and 2 data (n=19,629).

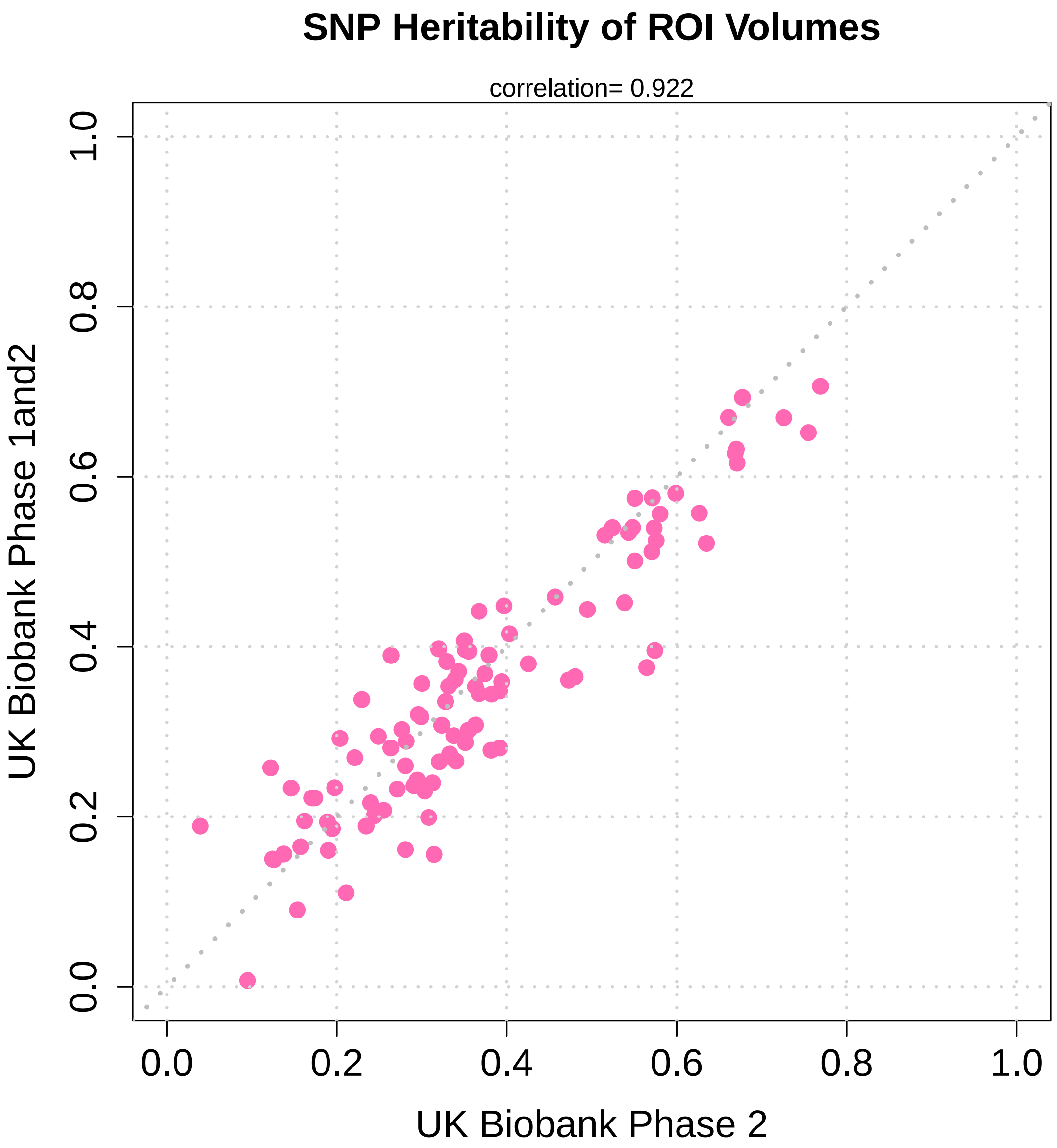


**Supplementary Figure 3:** Comparing SNP heritability estimates of ROI volumes in UK Biobank phase 2 data (n=10,431) and the combined phases 1 and 2 data (n=19,629).

**Supplementary Figure 4:** GWAS Manhattan and QQ plots for all the 101 ROI volumes, can be downloaded at `https://www.dropbox.com/s/0y3bsyeo9dcueky/ukbiobank_roi_volume_figures.zip?dl=0`.

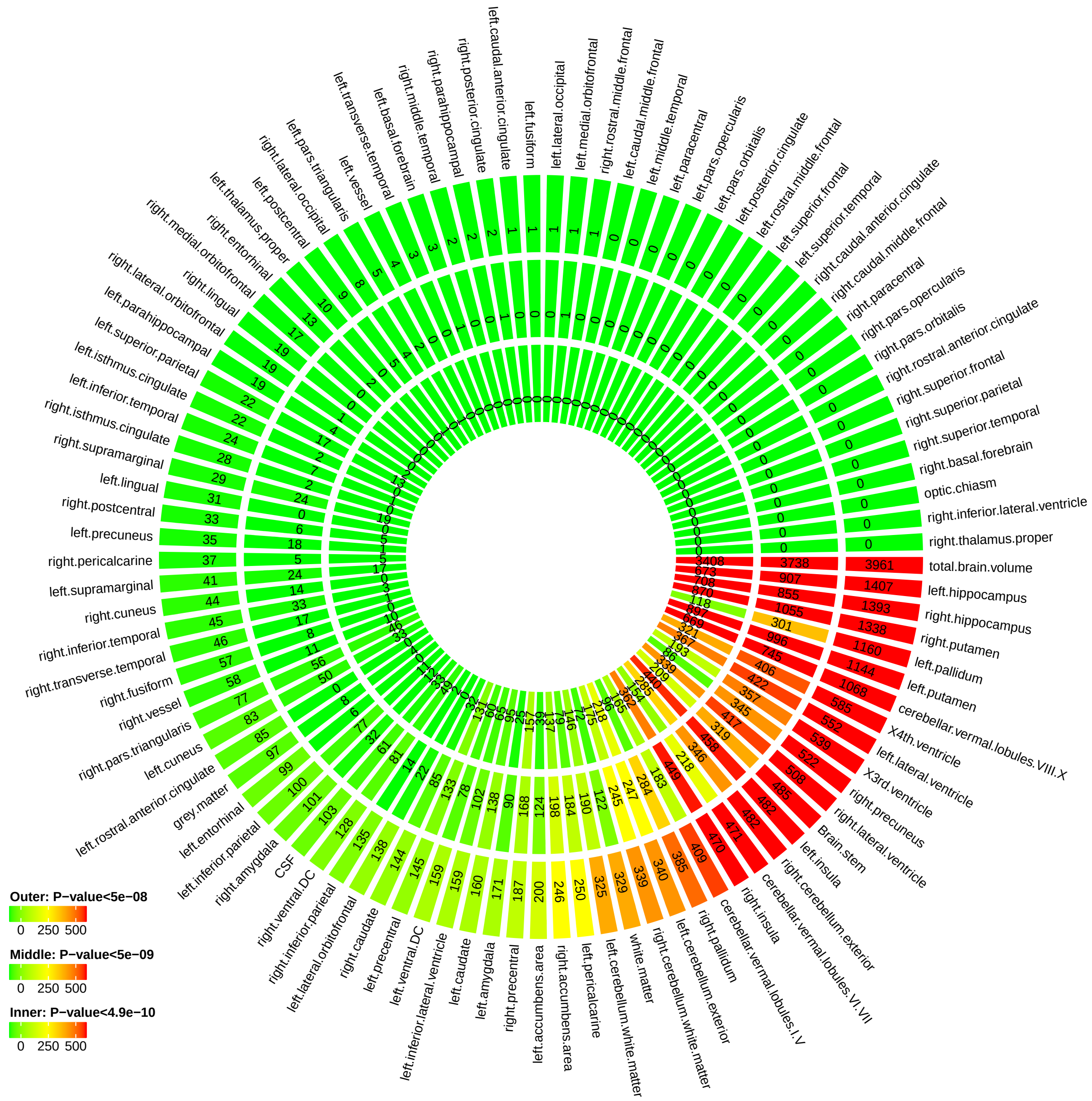


**Supplementary Figure 5:** Number of significant SNP associations discovered in UKB GWAS at different significance levels. Outer layer: P-value $< 5 \times 10^{-8}$; middle layer: P-value $< 5 \times 10^{-9}$; and inner layer: P-value $< 4.9 \times 10^{-10}$.

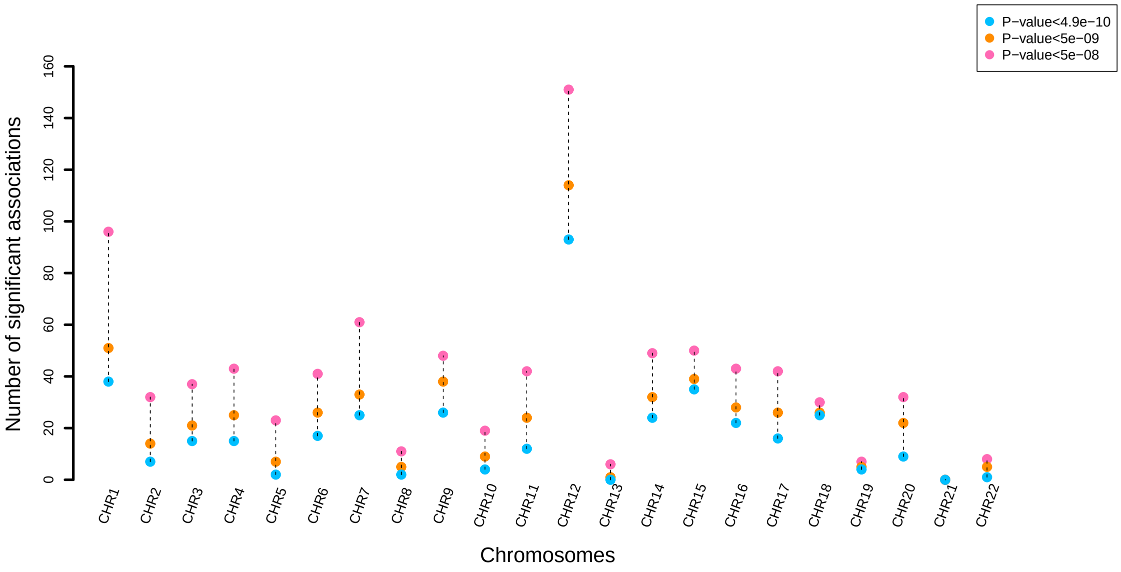


**Supplementary Figure 6:** Number of independent significant SNP associations discovered in UKB GWAS on each chromosome at different significance levels.

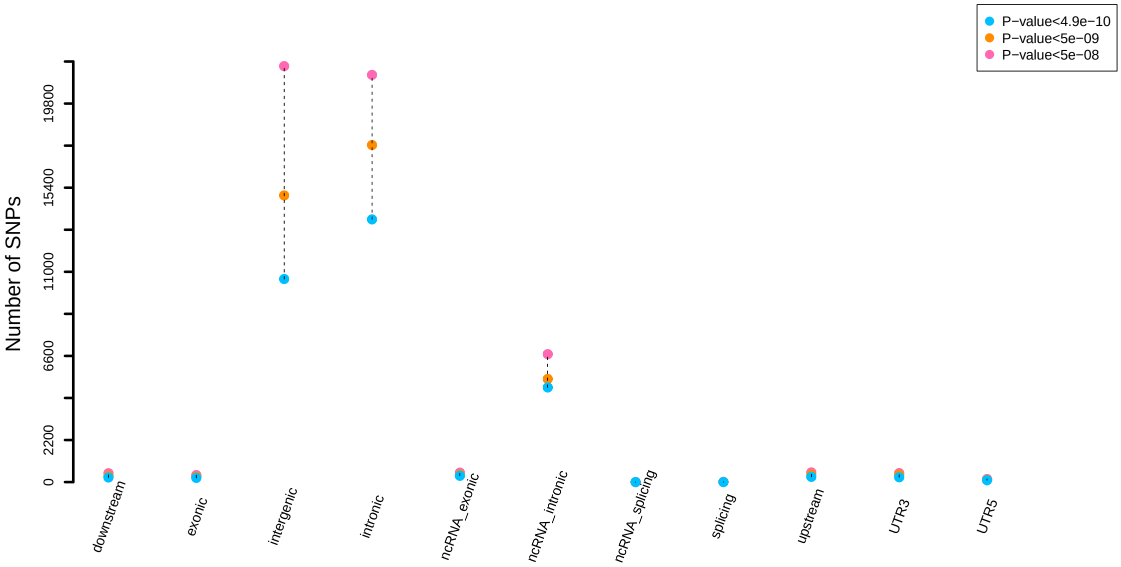


**Supplementary Figure 7:** Functional consequences of indepdenent SNPs (and SNPs in LD with them) indicated by functional annotation assigned by ANNOVAR at different significance levels.

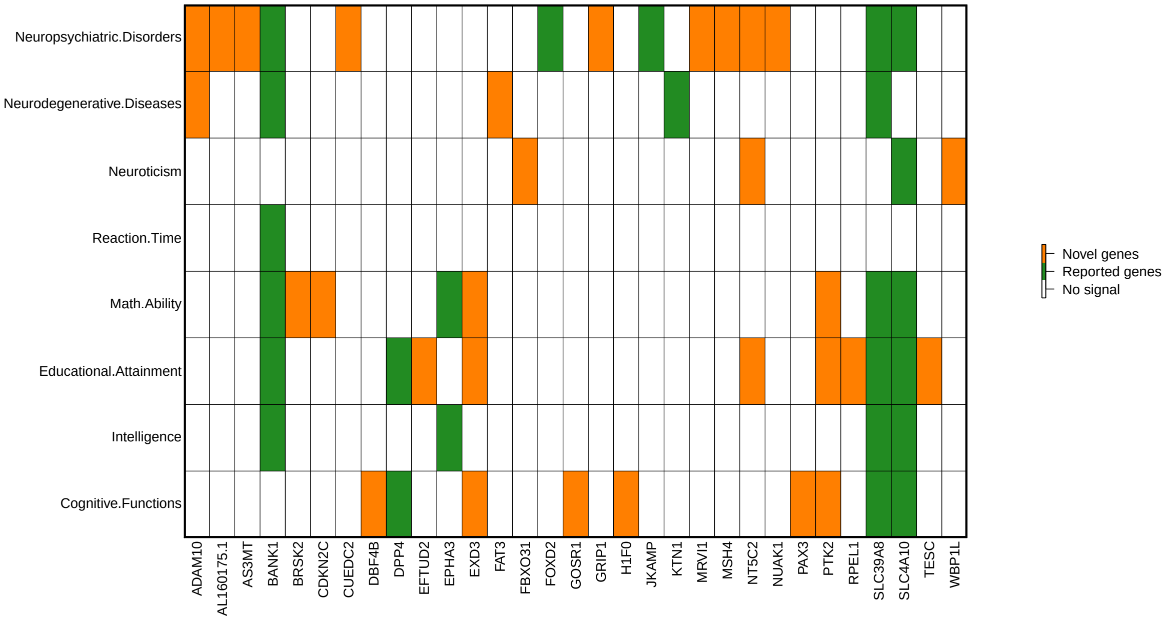


**Supplementary Figure 8:** Pleiotropic genes identified in functional mappings of ROI volumes that have been linked to cognitive traits and mental health disease/disorders in previous GWAS.

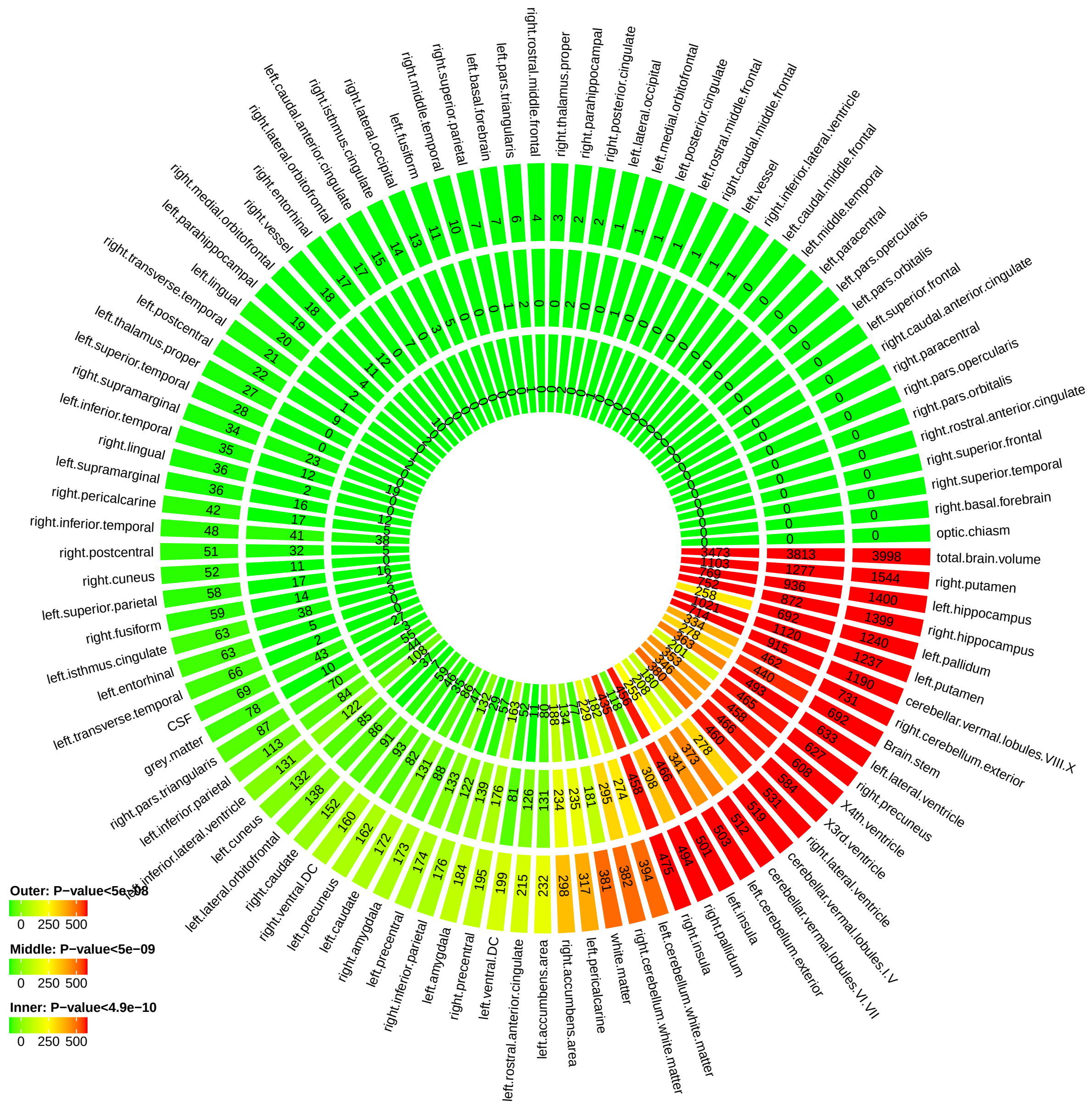


**Supplementary Figure 9:** Number of significant SNP associations discovered in meta-analyzed GWAS at different significance levels. Outer layer: P-value $< 5 \times 10^{-8}$; middle layer: P-value $< 5 \times 10^{-9}$; and inner layer: P-value $< 4.9 \times 10^{-10}$.

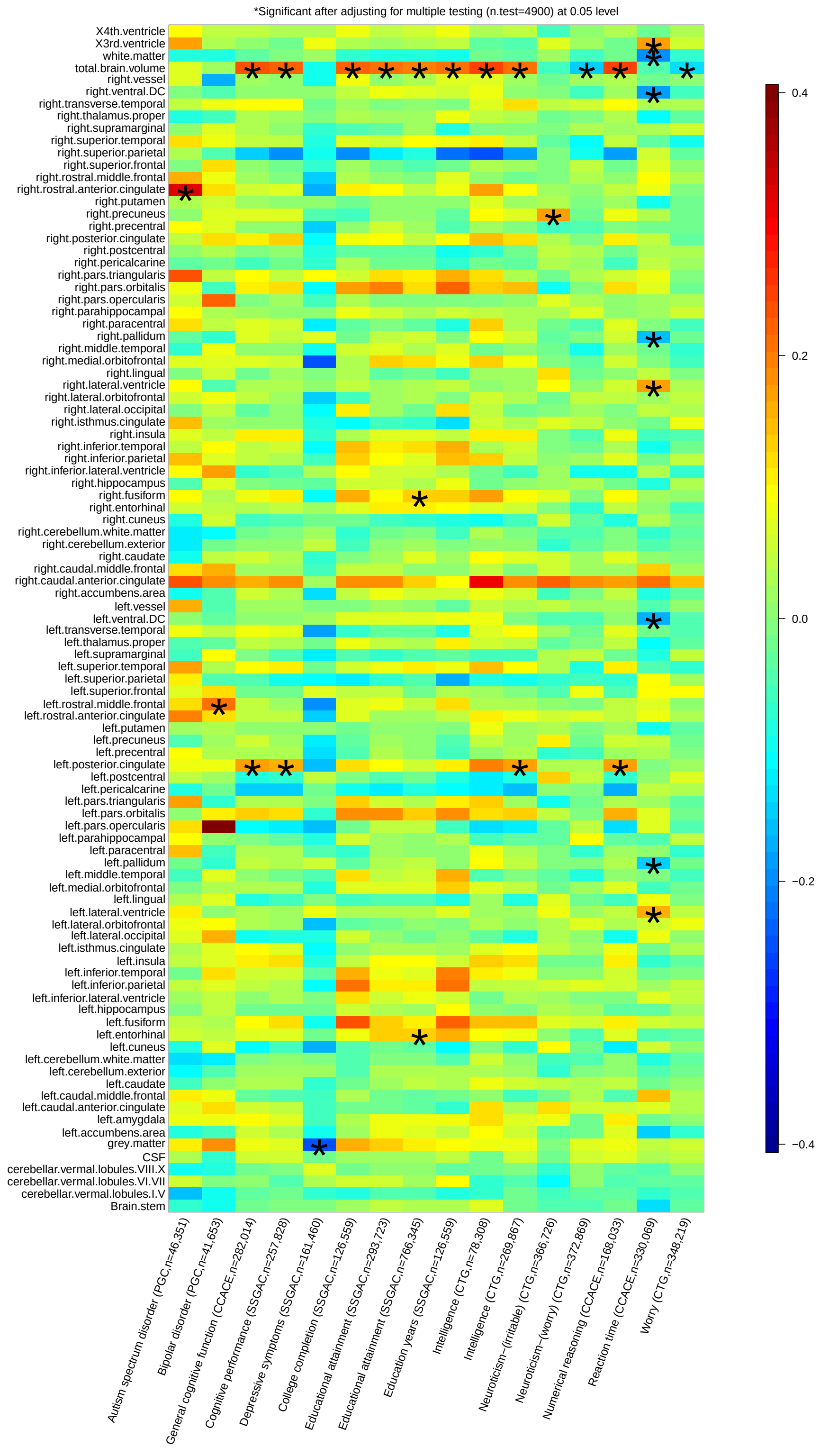


**Supplementary Figure 10:** Selected pairwise genetic correlations between ROI volumes and other traits. Stars are significant associations after adjusting for multiple testing by the Benjamini-Hochberg procedure at 0.05 level.

### 2 Supplementary tables

**Supplementary Table 1:** SNP heritability estimates of ROI volumes with using UKB phase 2 data (n=10,341) and UKB phases 1 and 2 data (n=19,629).

**Supplementary Table 2:** Number of significant SNP associations discovered in UKB GWAS at different significance levels.

**Supplementary Table 3:** List of independent significant SNP associations discovered in UKB GWAS at different significance levels.

**Supplementary Table 4:** Number of independent significant SNP associations discovered in UKB GWAS at different significance levels.

**Supplementary Table 5:** Number of independent significant SNP associations discovered in UKB GWAS on each chromosome at different significance levels.

**Supplementary Table 6:** List of significant genetic risk loci identified by UKB GWAS at different significance levels.

**Supplementary Table 7:** Number of significant genetic risk loci identified by UKB GWAS at different significance levels.

**Supplementary Table 8:** Number of significant genetic risk loci identified by UKB GWAS on each chromosome at different significance levels.

**Supplementary Table 9:** Summary of significant associations at different significance levels.

**Supplementary Table 10:** Independent significant (P-value $< 4.9 \times 10^{-10}$) SNPs and their correlated SNPs for ROI volumes that have previously been identified at P-value $< 9 \times 10^{-6}$ in GWAS of any traits listed in the NHGRI-EBI GWAS catalog (version 2019-01-31, www.ebi.ac.uk/gwas/).

**Supplementary Table 11:** Independent significant (P-value $< 4.9 \times 10^{-10}$) SNPs and their correlated SNPs for ROI volumes that have previously been identified at P-value $< 9 \times 10^{6}$ in GWAS of any brain volume and structure traits listed in the GWAS catalog (version 2019-01-31, www.ebi.ac.uk/gwas/).

**Supplementary Table 12:** Independent significant (P-value $< 5 \times 10^{-8}$) SNPs and their correlated SNPs for ROI volumes that have previously been identified at P-value $< 9 \times 10^{-6}$ in GWAS of any traits listed in the NHGRI-EBI GWAS catalog (version 2019-01-31, www.ebi.ac.uk/gwas/).

**Supplementary Table 13:** List of SNPs that are within LD of independent significant SNPs discovered by UKB GWAS at different significance levels.

**Supplementary Table 14:** List of significant gene-level associations identified by MAGMA (P-value $< 2 \times 10^{-8}$).

**Supplementary Table 15:** Functional consequences of indepdenent SNPs (and SNPs in LD with them) indicated by functional annotation assigned by ANNOVAR at different significance levels.

**Supplementary Table 16:** List of mapped genes identified in functional mapping of UKB GWAS results at different significance levels.

**Supplementary Table 17:** MAGMA gene-priority analysis for UKB GWAS results and 14 brain tissues. 14 brain tissues were from GTEx v7 RNA-seq database. Significant tissue groupings are highlighted in bold.

**Supplementary Table 18:** Number of significant SNP associations discovered in meta-analyzed GWAS at different significance levels.

**Supplementary Table 19:** Genetic correlations between several UKB ROIs volumes (TBV, left/right thalamus proper, left/right caudate, left/right putamen, left/right pallidum, left/right hippocampus, left/right accumbens area) and their corresponding traits studied in the ENIGMA consortium.

**Supplementary Table 20:** Sources of the 50 sets of publicly available GWAS summary statistics used in this study.

**Supplementary Table 21:** Genetic correlations estimates and p-values between ROI volumes and other traits.

**Supplementary Table 22:** Significant genetic correlation estimates and p-values between ROI volumes and other traits.

**Supplementary Table 23:** Prediction accuracy (incremental R-squared) of GWAS summary statistics using polygenic risk scores constructed on independent testing data.

**Supplementary Table 24:** Sample size and number of SNPs of the data used in each GWAS.

| Dataset | Sample Size | Number of SNPs |
|---:|---|---|
| UK Biobank | 19,629 | 8,944,375 |
| ADNI | 860 | 7,368,446 |
| HCP | 334 | 17,435,268 |
| PING | 461 | 8,612,247 |
| PNC | 537 | 5,151,137 |

**Supplementary Table 25:** Demographic information of five datasets.

| Dataset | Sample size | Mean age (s.d.) | Age range | Proportion of male |
|---:|---|---|---|---|
| UK Biobank | 19,629 | 62.51 (7.47) | (40 80) | 0.473 |
| ADNI | 1248 | 74.18 (7.16) | (55 92) | 0.566 |
| HCP | 1141 | 28.82 (3.68) | (22 36) | 0.542 |
| PING | 924 | 12.28 (4.99) | (3 21) | 0.518 |
| PNC | 1492 | 21.14 (3.71) | (14 29) | 0.515 |

### 3 Supplementary Note

#### Cohort information

In this study, we made use of data from five independent studies, whose demographic information is listed in Supplementary Table 25. The main GWAS was performed on the British individuals (self-reported ethnic background, Data-Field 21000) in UK Biobank study. For the other four cohorts, we only considered the unrelated European ancestry individuals (self-reported race, ethnic and family information) in GWAS. In polygenic risk score prediction, we used all available individuals (with SNP and phenotype data) to examine the prediction power of UKB GWAS results in testing data.

#### Genotyping and quality control

We downloaded the imputed SNP data from UKB and HCP data resources, respectively. Genotype imputation was performed locally on the PNC, ADNI, and PING datasets using consistent procedures via MACH-Admix (Liu et al., 2013). A full description of the imputation procedures in PNC, ADNI, and PING datasets was detailed supplementary information of Zhao et al. (2017). We further performed the following SNP data quality controls on each dataset: 1) exclude subjects with more than 10% missing genotypes; 2) exclude SNPs with minor allele frequency less than 0.01; 3) exclude SNPs with larger than 10% missing genotyping rate; 4) exclude markers that failure the Hardy-Weinberg test at $1 \times 10^{-7}$ level; and 5) remove SNPs with imputation INFO score less than 0.8.

#### PING Methods

Part of the data used in the preparation of this article were obtained from the Pediatric Imaging, Neurocognition and Genetics (PING) Study database (`http://ping.chd.ucsd.edu/`). PING was launched in 2009 by the National Institute on Drug Abuse (NIDA) and the Eunice Kennedy Shriver National Institute Of Child Health & Human Development (NICHD) as a 2-year project of the American Recovery and Reinvestment Act. The primary goal of PING has been to create a data resource of highly standardized and carefully curated magnetic resonance imaging (MRI) data, comprehensive genotyping data, and developmental and neuropsychological assessments for a large cohort of developing children aged 3 to 20 years. The scientific aim of the project is, by openly sharing these data, to amplify the power and productivity of investigations of healthy and disordered development in children, and to increase understanding of the origins of variation in neurobehavioral phenotypes. For up-to-date information, see `http://ping.chd.ucsd.edu/`.

### ADNI Methods

Data used in the preparation of this article were obtained from the Alzheimers Disease Neuroimaging Initiative (ADNI) database (`http://adni.loni.usc.edu`). The ADNI was launched in 2003 by the National Institute on Aging (NIA), the National Institute of Biomedical Imaging and Bioengineering (NIBIB), the Food and Drug Administration (FDA), private pharmaceutical companies and non-profit organizations, as a 60 million, 5-year public-private partnership. The primary goal of ADNI has been to test whether serial magnetic resonance imaging (MRI), positron emission tomography (PET), other biological markers, and clinical and neuropsychological assessment can be combined to measure the progression of mild cognitive impairment (MCI) and early Alzheimers disease (AD). Determination of sensitive and specific markers of very early AD progression is intended to aid researchers and clinicians to develop new treatments and monitor their effectiveness, as well as lessen the time and cost of clinical trials.

The Principal Investigator of this initiative is Michael W. Weiner, MD, VA Medical Center and University of California San Francisco. ADNI is the result of efforts of many co-investigators from a broad range of academic institutions and private corporations, and subjects have been recruited from over 50 sites across the U.S. and Canada. The initial goal of ADNI was to recruit 800 subjects but ADNI has been followed by ADNI-GO and ADNI-2. To date these three protocols have recruited over 1500 adults, ages 55 to 90, to participate in the research, consisting of cognitively normal older individuals, people with early or late MCI, and people with early AD. The follow up duration of each group is specified in the protocols for ADNI-1, ADNI-2 and ADNI-GO. Subjects originally recruited for ADNI-1 and ADNI-GO had the option to be followed in ADNI-2. For up-to-date information, see `www.adni-info.org`.

### Pediatric Imaging, Neurocognition and Genetics (PING) Authors

Connor McCabe[1], Linda Chang[2], Natacha Akshoomoff[3], Erik Newman[1], Thomas Ernst[2], Peter Van Zijl[4], Joshua Kuperman[5], Sarah Murray[6], Cinnamon Bloss[6], Mark Appelbaum[1], Anthony Gamst[1], Wesley Thompson[3], Hauke Bartsch[5].

### Alzheimer's Disease Neuroimaging Initiative (ADNI) Authors

Michael Weiner[7], Paul Aisen[1], Ronald Petersen[8], Clifford R. Jack Jr[8], William Jagust[9], John Q. Trojanowki[10], Arthur W. Toga[11], Laurel Beckett[12], Robert C. Green[13], Andrew

J. Saykin[14], John Morris[15], Leslie M. Shaw[10], Zaven Khachaturian[16], Greg Sorensen[17], Maria Carrillo[18], Lew Kuller[19], Marc Raichle[15], Steven Paul[20], Peter Davies[21], Howard Fillit[22], Franz Hefti[23], Davie Holtzman[15], M. Marcel Mesulman[24], William Potter[25], Peter J. Snyder[26], Adam Schwartz[27], Tom Montine[28], Ronald G. Thomas[1], Michael Donohue[1], Sarah Walter[1], Devon Gessert[1], Tamie Sather[1], Gus Jiminez[1], Danielle Harvey[12], Matthew Bernstein[8], Nick Fox[29], Paul Thompson[11], Norbert Schuff[7], Charles DeCarli[12], Bret Borowski[8], Jeff Gunter[8], Matt Senjem[8], Prashanthi Vemuri[8], David Jones[8], Kejal Kantarci[8], Chad Ward[8], Robert A. Koeppe[30], Norm Foster[31], Eric M. Reiman[32], Kewei Chen[32], Chet Mathis[19], Susan Landau[9], Nigel J. Cairns[15], Erin Householder[15], Lisa Taylor-Reinwald[15], Virginia M.Y. Lee[10], Magdalena Korecka[10], Michal Figurski[10], Karen Crawford[11], Scott Neu[11], Tatiana M. Foroud[14], Steven Potkin[33], Li Shen[14], Kelley Faber[14], Sungeun Kim[14], Kwangsik Nho[14], Leon Thal[1], Richard Frank[34], Neil Buckholtz[35], Marilyn Albert[36], John Hsiao[35].

[1]UC San Diego, La Jolla, CA 92093, USA. [2]U Hawaii, Honolulu, HI 96822, USA. [3]Department of Psychiatry, University of California, San Diego, La Jolla, California 92093, USA. [4]Kennedy Krieger Institute, Baltimore, MD 21205, USA. [5]Multimodal Imaging Laboratory, Department of Radiology, University of California San Diego, La Jolla, California 92037, USA. [6]Scripps Translational Science Institute, La Jolla, CA 92037, USA. [7]UC San Francisco, San Francisco, CA 94143, USA. [8]Mayo Clinic, Rochester, MN 55905, USA. [9]UC Berkeley, Berkeley, CA 94720-5800, USA. [10]U Pennsylvania, Philadelphia, PA 19104, USA. [11]USC, University of Southern California, Los Angeles, CA 90033, USA. [12]UC Davis, Davis, CA 95616, USA. [13]Brigham and Women s Hospital/Harvard Medical School, Boston MA 02115, USA. [14]Indiana University, Indianapolis, IN 46202-5143, USA. [15]Washington University St. Louis, St. Louis, MO 63130, USA. [16]Prevent Alzheimers Disease 2020, Rockville, MD 20850, USA. [17]Siemens [18]Alzheimers Association, Chicago, IL 60601, USA. [19]University of Pittsburgh, Pittsburgh, PA 15260, USA. [20]Cornell University, Ithaca, NY 14850, USA. [21]Albert Einstein College of Medicine of Yeshiva University, Bronx, NY 10461, USA. [22]AD Drug Discovery Foundation, New York, NY 10019, USA. [23]Acumen Pharmaceuticals, Livermore, California 94551, USA. [24]Northwestern University, Evanston, IL 60208, USA. [25]National Institute of Mental Health, Bethesda, MD 20892-9663, USA. [26]Brown University, Providence, RI 02912, USA. [27]Eli Lilly, Indianapolis, Indiana 46285, USA. [28]University of Washington, Seattle, WA 98195, USA. [29]University of London, London WC1E 7HU, UK. [30]University of Michigan, Ann Arbor, MI 48109, USA. [31]University of Utah, Salt Lake City, UT 84112, USA. [32]Banner Alzheimers Institute, Phoenix, AZ 85006, USA. [33]UC Irvine, Irvine, CA 92697, USA. [34]General Electric [35]National Institute on Aging/National Institutes of Health, Bethesda, MD 20892, USA. [36]The Johns Hopkins University, Baltimore, MD 21218, USA.

#### LD Hub

We gratefully acknowledge all the studies and databases that made GWAS summary data available: ADIPOGen (Adiponectin genetics consortium), C4D (Coronary Artery Disease Genetics Consortium), CARDIoGRAM (Coronary ARtery DIsease Genome wide Replication and Meta-analysis), CKDGen (Chronic Kidney Disease Genetics consortium), dbGAP (database of Genotypes and Phenotypes), DIAGRAM (DIAbetes Genetics Replication And Meta-analysis), ENIGMA (Enhancing Neuro Imaging Genetics through Meta Analysis), EAGLE (EArly Genetics & Lifecourse Epidemiology Eczema Consortium, excluding 23andMe), EGG (Early Growth Genetics Consortium), GABRIEL (A Multidisciplinary Study to Identify the Genetic and Environmental Causes of Asthma in the European Community), GCAN (Genetic Consortium for Anorexia Nervosa), GEFOS (GEnetic Factors for OSteoporosis Consortium), GIANT (Genetic Investigation of ANthropometric Traits), GIS (Genetics of Iron Status consortium ), GLGC (Global Lipids Genetics Consortium), GPC (Genetics of Personality Consortium), GUGC (Global Urate and Gout consortium), HaemGen (haemotological and platelet traits genetics consortium), HRgene (Heart Rate consortium), IIBDGC (International Inflammatory Bowel Disease Genetics Consortium), ILCCO (International Lung Cancer Consortium), IMSGC (International Multiple Sclerosis Genetic Consortium), MAGIC (Meta-Analyses of Glucose and Insulin-related traits Consortium), MESA (Multi-Ethnic Study of Atherosclerosis), PGC (Psychiatric Genomics Consortium), Project MinE consortium, ReproGen (Reproductive Genetics Consortium), SSGAC (Social Science Genetics Association Consortium) and TAG (Tobacco and Genetics Consortium), TRICL (Transdisciplinary Research in Cancer of the Lung consortium), UK Biobank. We gratefully acknowledge the contributions of Alkes Price (the systemic lupus erythematosus GWAS and primary biliary cirrhosis GWAS) and Johannes Kettunen (lipids metabolites GWAS).
